# ORMDL3-mediated SPT regulation Coordinates Myelin Sphingolipid and Protein Synthesis in Oligodendrocytes

**DOI:** 10.64898/2026.09.23.753788

**Authors:** Usha Mahawar, Raheema Fulani, Karli Mockenhaupt, Sumit Saha, Faheem Farooq, Fatemah S. Afshari, Susmita Manam, Carmen Sato-Bigbee, Babette Fuss, Jeffrey Dupree, Tomasz Kordula, Brian W Wattenberg

## Abstract

Myelin is an essential and highly specialized membrane in the central and peripheral nervous systems that enwraps axons to accelerate electrical transmission and support neuronal health. The generation of this multilamellar structure requires a tightly coordinated synthesis of specific proteins and lipids. Among these are sphingolipids (SLs), which are major components of myelin. SL production is initiated by the serine palmitoyltransferase (SPT) enzyme complex, the rate-limiting enzyme in the *de novo* SL biosynthesis pathway. ORMDL proteins (ORMDL1-3) are the regulatory subunits of SPT, which, by sensing ceramide levels, tune SL flux. Although ORMDL3 has been linked to asthma and peripheral myelination, its role in CNS myelination and the CNS myelin-making oligodendrocytes (OLGs) remains unclear. We therefore assessed the function of ORMDL3 in OLs by generating a *Cnp-Cre*-driven, oligodendrocyte-specific *Ormdl3* conditional knockout (cKO) mouse model. Loss of *Ormdl3* selectively increased myelin SLs, particularly long-chain sulfatides, without altering ceramide or galactosylceramide abundance. These changes are most prominent around postnatal day 35, a developmental period of active myelin turnover. Ultrastructural analysis of optic nerves shows a thicker myelin sheath and increased axon caliber in cKO mice. Unexpectedly, deletion of *Ormdl3* also increases levels of major myelin proteins (MBP, MOG, and PLP) and is accompanied by dynamic, region- and sex-dependent regulation of enzymes involved in sulfatide biosynthesis, without altering OLGs number or maturation. Together, for the first time, these findings identify *Ormdl3* as a key regulator of SL homeostasis during developmental myelination and suggest that it helps synchronize lipid synthesis with myelin protein expression.

## 1. Introduction

The CNS relies on coordinated communication between neurons and glial cells to process and transmit information [1]. Among glia, oligodendrocytes (OLGs) generate myelin, a multilamellar sheath that insulates axons to enable rapid action potential propagation [2] and provides metabolic support to the underlying neuron [3]. Myelin is a specialized lipid-rich membrane composed of ∼70-80% lipids (by dry weight) and ∼20-30% proteins, including, among others, myelin basic protein (MBP), myelin oligodendrocyte glycoprotein (MOG), and proteolipid protein (PLP) [4]. During early CNS myelin development, lipids and proteins are produced in a coordinated sequence, with lipid synthesis preceding myelin protein production [5, 6]. Once lipids are produced, OLGs produce proteins that help assemble and stabilize a multilayered sheath around axons [7]. Myelin proteins, such as MBP, promote adhesion between compacted layers, PLP supports mature sheath architecture and axon–glia metabolic coupling, and MOG contributes to outer-sheath integrity and communication with other cell types [4, 8]. Disruption in myelin production contributes to congenital dysmyelination, acquired demyelination, and immune-mediated demyelination, including Krabbe’s disease, leukodystrophy, and multiple sclerosis, which impair nerve conduction and can drive progressive cognitive and motor decline [9, 10]. Sphingolipids (SLs), cholesterol, and phospholipids are the major constituents of myelin lipids that provide structural stability and functional properties [11]. Among them, SL contributes to membrane organization, signaling, and lipid-raft formation, which helps organize myelin proteins [12–14]. Galactosylceramides, sulfatides, and sphingomyelin are the predominant SL species in myelin [5, 15]. The *de novo* biosynthesis of SL is tightly regulated, with serine palmitoyltransferase (SPT) catalyzing the first and rate-limiting step of the SL biosynthesis pathway **(Fig. 1A)** [16]. SPT is an endoplasmic reticulum (ER) bound multi-subunit enzyme complex. Previous studies from our lab in the developing rat brain have shown that SPT subunit composition at the mRNA level, including *Sptlc1, Sptlc2, Sptlc3*, and the small subunits *Sptssa* and *Sptssb*, changes dynamically during OLG maturation and myelination, possibly affecting the types of SL produced [6]. SPTLC1 anchors the serine palmitoyltransferase complex to the endoplasmic reticulum and associates with either SPTLC2 or SPTLC3 to form the catalytic core [16]. The specific composition of this core determines substrate preference and, therefore, the chain length of the sphingoid base produced [17]. The canonical SPTLC1/SPTLC2/SPTssa complex preferentially condenses serine with palmitoyl-CoA to generate 18-carbon sphingoid bases. In contrast, incorporation of SPTLC3 enables the utilization of shorter acyl-CoA substrates, such as myristoyl-CoA, leading to the production of shorter sphingoid bases and the formation of non-canonical SL species. The inclusion of the SPTssb subunit further expands substrate flexibility, allowing SPT to accept both shorter and longer acyl-CoAs and thereby generate a diverse spectrum of sphingoid bases with varying chain lengths. Thus, changes in SPT subunit expression or assembly can alter both the amount and composition of SL produced.

**Figure 1:**
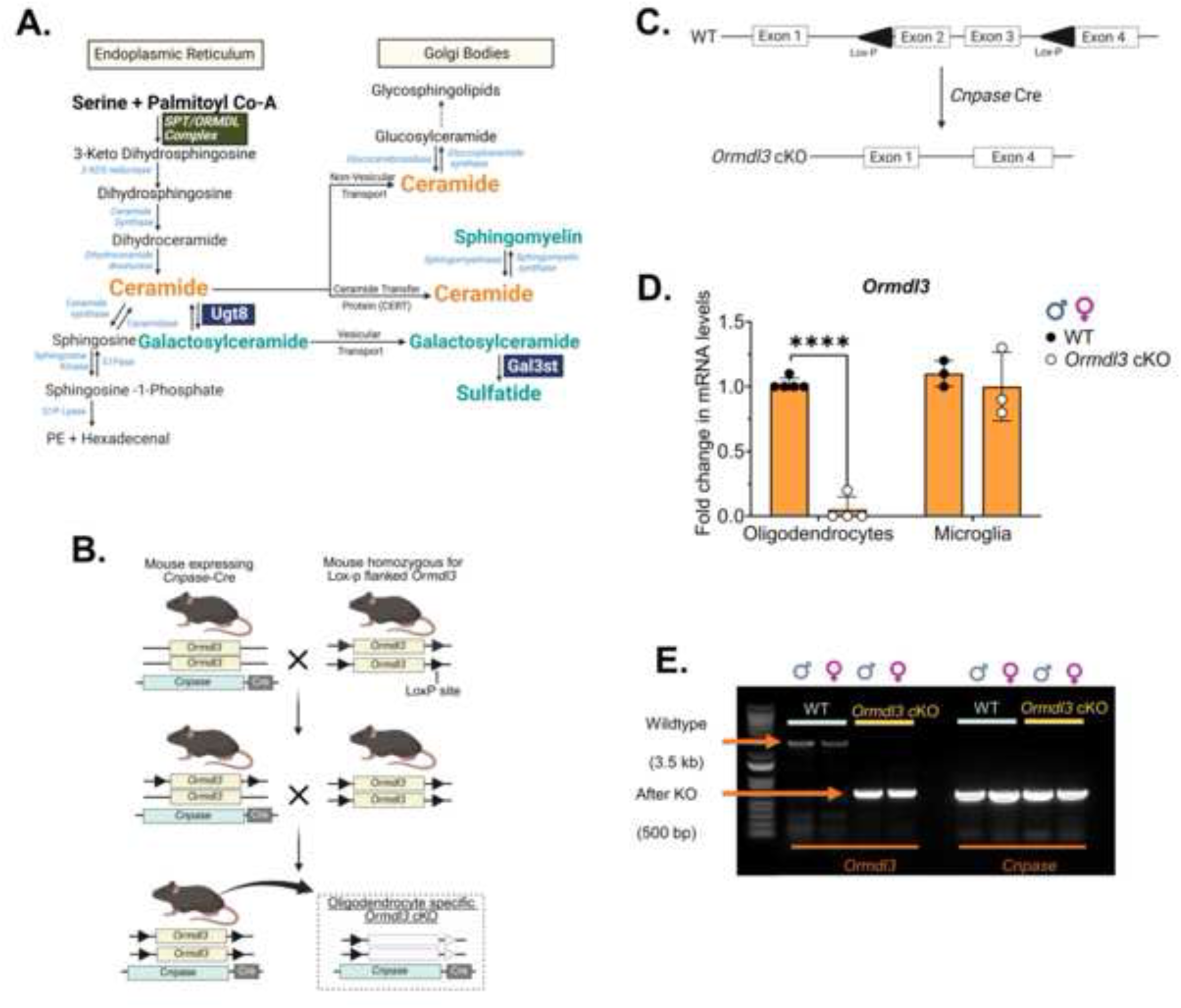
Successful verification of Oligodendrocyte-specific *Ormdl3* cKO in the brain of postnatal day 16 mice. **A.)** Cartoon representation of the *de novo* sphingolipid biosynthesis pathway. The major sphingolipid species present in myelin are highlighted in teal green color. **B.)** Schematic representation of the generation of oligodendrocyte-specific conditional *Ormdl3* knockout (cKO) mice. The *Cnp*-Cre mice are shown in a bluish-green color. Mice carrying loxP sites around the axon of *Ormdl3* are shown in yellow boxes. **C.)** Schematic illustration showing loxP sites (black triangles) at the beginning of exon 2 and exon 4 of *Ormdl3*. Cre excises exon 2, exon 3, and a part of exon 4 of *Ormdl3*, resulting in the generation of an *Ormdl3* cKO. **D.)** mRNA levels of *Ormdl3* were measured in isolated oligodendrocytes and microglia from postnatal day 16 WT and *Ormdl3* cKO mice. Data are shown as the mean ± SD; n = 3–5. Statistical significance was tested with the Student’s two-tailed t-test. *Asterisks* denote significance, p <0.00005 = ****. **E.)** PCR amplification of *Ormdl3* and CNP in total genomic DNA was performed using brain lysates extracted from postnatal day 35 WT and *Ormdl3* cKO mice. CNP: 2’,3’-Cyclic nucleotide 3’-phosphohydrolase; PCR: polymerase chain reaction; cKO: conditional KO.

SPT is homeostatically regulated by its regulatory subunits, the ORMDLs (ORMDL1, ORMDL2, and ORMDL3). The ORMDLs directly sense cellular ceramide levels through a specific ceramide binding site, which, when occupied, stabilizes the ORMDL amino terminus into the catalytic site of SPT [18, 19]. Each ORMDL isoform senses different ceramide levels to regulate SPT [18]. While the roles of ORMDLs in cancer [20], immune function [21], and asthma [22, 23] have been explored, their specific contributions to OLG biology and CNS myelination remain incompletely understood. Notably, whole-body *Ormdl* knockout mouse studies show that loss of ORMDL proteins causes severe dysmyelination in the peripheral nervous system (PNS) [24]. In addition, global deletion of all three *Ormdl* isoforms is embryonically lethal [24]. Whole-body *Ormdl1/3* double-knockout mice exhibit redundant myelin in the sciatic nerve at 8 weeks of age, suggesting that SL biosynthesis regulation can affect myelin structure and function in the PNS. Consistently, overexpression of SPTLC1/SPTLC2/SPTSSA fusion constructs in Schwann cells phenocopied the *Ormdl1/3* double knockout by increasing SL production, highlighting the importance of SL homeostasis by SPT regulation for proper PNS myelination.

In the field of OLG biology, much attention has focused on OLG differentiation [25–27], the importance of the myelin proteins[4, 28–30], mechanisms by which OLGs and neurons communicate during myelination[31], and the contributions of astrocytes and microglia to myelination [32]. Additionally, the mechanisms by which OLGs produce cholesterol and the role of cholesterol in myelin have been studied [33, 34]. Furthermore, efforts have been made to investigate the roles of various enzymes involved in SL biosynthesis, including sphingomyelin [34] and sulfatides [35, 36].

Despite these advances, several key questions in myelin SL biology remain unanswered. In particular, the cell–type–specific mechanisms that regulate the production of myelin-associated SLs are not well understood. It is also unclear how *Ormdl*-mediated regulation of SPT influences the synthesis of distinct SL species during CNS development. Dynamic changes in *Ormdl* isoform expression at the mRNA level during rat brain OLG development further suggest that different *Ormdl* isoforms may regulate SL biosynthesis in a temporally and spatially specific manner[6].

The central question we are addressing here is how OLG-specific regulation of SL biosynthesis by ORMDL3 influences CNS myelination. We focused on *Ormdl3* because its expression changes dynamically during myelination and, among *Ormdl* isoforms, it is the most sensitive to ceramide levels [6, 18].

To test this, we generated an OLG-specific *Ormdl3* conditional knockout (cKO) using Cre-lox recombination driven by the *Cnp* promoter, which is active during early brain development and shows peak enzyme activity before postnatal day 10 [37]. To assess the impact of *Ormdl3* cKO in the CNS, we examined the optic nerves and the brain. The knockout markedly altered SL levels, ultrastructure, and, unexpectedly, myelin protein abundance. Collectively, these findings provide compelling evidence that ORMDL3 is an essential regulator of myelin biogenesis in OLGs, primarily by modulating SPT activity and SL production. Increased SL production, in turn, coordinately elevates myelin protein abundance through an as-yet-unknown mechanism.

## 2. Materials and Methods

### 2.1. Materials

#### 2.1.1. Immunoblotting

The following antibodies were used in this study: anti-Myelin Basic Protein (MBP) (EMD Millipore #MAB386), anti-Myelin Oligodendrocyte Glycoprotein (MOG) (EMD Millipore #MAB5680), anti-proteolipid protein (PLP) (Abcam #ab28486), anti-β-actin (Invitrogen #PA1-16889), anti-UDP-glycosyltransferase 8 (UGT8) (Proteintech #17982-1-AP), anti-galactose-3-O-sulfotransferase 1 (GAL3ST1) (Proteintech #24851-1-AP), anti-Calnexin (ENZO #ADI-SPA-860-D), anti-SPTLC1 (anti-LCB1) (BD Transduction laboratories #611305), anti-glyceraldehyde-3-phosphate-dehydrogenase (GAPDH) (Invitrogen #MA5-35235).

HRP-conjugated secondary antibodies for a mouse (Thermo Fisher #31430), rabbit (Thermo Fisher #31460), and goat anti-rat (Jackson Immuno Research Laboratories #112-035-003). Amersham ECL Prime immunoblot detection reagent (Thermo Fisher #RPN2322), 0.22 µm PVDF membrane (Bio-Rad), blotting grade blocker (Bio-Rad #1706404), and Fat-free BSA (Fisher #BP9704100)

#### 2.1.2. RNA isolation and quantitative real-time PCR reagents

Trizol reagent (Thermo Fisher #15596026), 5-prime heavy 2 ml-phase lock gel tubes (VWR #2302830), high-capacity reverse transcriptase kit ( Applied Biosystems #4368814), thin-wall 96-well plates (Biorad #HSP9601), micro seal optical plate covers (Biorad #MSB1001), GlycoBlue precipitate (Thermo # AM9515), TaqMan FAM-labelled PrimeTime qPCR probes (Integrated DNA Technologies) (details in **supplementary table 4**), PrimeTime gene expression master mix ( IDT #1055772), and SYBR Green qPCR master mix (Biorad) (details in **supplementary table 3 and 5**).

#### 2.1.3. Chemicals and reagents

Percoll (GE Healthcare Bio-Science #17-0891-01), HBSS (Gibco #14185-052), DMEM/Ham’s F21 medium (Gibco #11320-033), HEPES (Sigma-Aldrich #H7006-500G), bovine pancreas DNase (Sigma-Aldrich #DN-25), papain (Sigma-Aldrich #P-4762), glucose (Sigma-Aldrich #G5146), sucrose (Sigma-Aldrich #57-50-1), lead citrate (Sigma-Aldrich #15326), trizma hydrochloride (Sigma-Aldrich T3253-1KG), chloroform/isoamyl alcohol (ThermoFisher Scientific #15593031), EDTA-free protease inhibitor (ThermoFisher Scientific #A32965), Millonigs Buffer (Electron Microscopy Sciences #11582-01), EM grade aqueous glutaraldehyde (Electron Microscopy Sciences #16310), paraformaldehyde(Electron Microscopy Sciences #15700), sodium cacodylate (Electron Microscopy Sciences #12300), osmium tetroxide (Polysciences), propylene oxide (Polysciences #00236), PolyBed 812 resin (Polysciences #08791), dodencylsuccinic anhydride (Polysciences #00563), nadic methyl anhydride (Polysciences #00886), DMP-30 (Polysciences #00553), uranyl acetate (Polysciences #21447-25), sodium hydroxide (VWR #1310-73-2), sodium acetate (VWR #6131-90-4), isopropanol (VWR #BP2618500), EDTA (VWR #6381-92-6), sodium bicarbonate (VWR #S233-500), potassium chloride (VWR #P217-500), magnesium chloride hexahydrate (VWR #M33-500), sodium chloride (VWR #BP358-212), magnesium sulfate heptahydrate (VWR #2500-01), sodium phosphate monobasic monohydrate (VWR #4011-01), calcium chloride (ICN biomedicals).

#### 2.1.4. RNAscope and immunohistochemistry

2-2-2 tribromoethanol (Sigma-Aldrich #T48402), Tissue-Tek O.C.T. compound (Sakura Finetek #4583), Leica Bond Plus slides (Leica Biosystems # S21.2113.A), RNAscope 2.5 LS Probe probes (Advanced Cell Diagnostics): Mm-*Ormdl3*-C1 (#445338-C1), Mm-*Olig2*-C2 (# 447098-C2), Mm-*Plp1*-C4 (# 428188-C4); for fluorescent detection, RNAscope LS Fluorescent Reagent and RNAscope LS 4-Plex Ancilliary kits (Advanced Cell Diagnostics #322800 and #322830), Opal fluorophores (Akoya Biosciences): Opal 690 (#PN FP1497001KT), Opal 620 (# PN FP1495001KT), Opal 520 (# PN FP1487001KT), The following antibodies were used for immunohistochemistry: anti-Olig2 (Cat. No OLIG2-0100, Aves Labs; RRID: AB_2924438), CC1 (Cat. No. OP80, Millipore, RRID: AB_2057371), goat anti-Chicken IgY, Alexa Fluor 633 (Cat No. A-21103, Thermo Fisher Scientific, RRID: AB_2535756), and goat anti-Mouse IgG2b, Alexa Fluor 568 (Cat No. A21144, Thermo Fisher Scientific, RRID: AB_2535780). Vectashield (Cat. No. H-1500-10, Vector Laboratories, Burlingame, CA), using Hoechst 33342 (Cat. No. 14533; Millipore Sigma, Burlington, MA)

#### 2.1.5. Lipidomics

Internal standards were purchased from Avanti Polar Lipids (Alabaster, AL). Internal standards were added to samples in a 10 µl ethanol:methanol:water (7:2:1) cocktail at 250 pmol each. Standards for sphingoid bases and sphingoid base 1-phosphates were 17-carbon chain length analogues: C17-sphingosine, (2S,3R,4E)-2-aminoheptadec-4-ene-1,3-diol (d17:1-So); C17-sphinganine, (2S,3R)-2-aminoheptadecane-1,3-diol (d17:0-Sa); C17-sphingosine 1-phosphate, heptadecasphing-4-enine-1-phosphate (d17:1-So1P); and C17-sphinganine 1-phosphate, heptadecasphinganine-1-phosphate (d17:0-Sa1P). Standards for N-acyl sphingolipids were C12-fatty acid analogs: C12-Cer, N-(dodecanoyl)-sphing-4-enine (d18:1/C12:0); C12-lactosylceramide, N-(dodecanoyl)-1-β-lactosyl-sphing-4-eine (d18:1/C12:0 LacCer); C12-sphingomyelin, N-(dodecanoyl)-sphing-4-enine-1-phosphocholine (d18:1/C12:0-SM); and C12-glucosylceramide, N-(dodecanoyl)-1-β-glucosyl-sphing-4-eine (d18:1/C12:0 GlcCer). For LC-MS/MS analyses, a Shimadzu Nexera LC-30 AD binary pump system coupled to a SIL-30AC autoinjector and a DGU20A5R degasser, and an AB Sciex 5500 quadrupole/linear ion trap (QTrap) (SCIEX Framingham, MA) operating in triple quadrupole mode, was used. Q1 and Q3 were set to pass molecularly distinctive precursor and product ions (or a scan across multiple m/z values in Q1 or Q3), using N2 to collisionally induce dissociation in Q2 (which was offset from Q1 by 30-120 eV); the ion source temperature was set to 500 °C. Teflon-lined cap (VWR #60827-453)

### 2.2. Methods

#### 2.2.1. Generation of the oligodendrocyte-specific *Ormdl3* conditional KO mice model

The *Ormdl3* floxed mice, which contain LoxP sites, were obtained from the Jackson Laboratory (B6.Cg-*Ormdl3*tm1.1Rlp/J). These mice were then crossed with *Cnp*-Cre mice, provided by Dr. Tomasz Kordula at VCU, to produce OLG-specific *Ormdl3* cKO mice. The *Cnp*-Cre mouse line was originally developed and characterized by Dr. Klaus-Armin Nave at the Max Planck Institute in Germany [38]. The mouse line was maintained at Virginia Commonwealth University in accordance with the Institutional Animal Care and Use Committee guidelines. Mice in animal facilities were kept in a 12-hour light/dark cycle, with standard laboratory breeder’s chow and water. Littermates were separated into male and female groups for all experiments.

#### 2.2.2. Isolation of primary OLGs from the mouse brain

Primary OLGs were isolated from 16 and 30-day old mice by following a Percoll gradient centrifugation protocol [39]. Animals were anesthetized using isoflurane for 30 seconds, followed by rapid decapitation. The cerebral hemispheres were rapidly dissected out, and after removal of the meninges, the tissue corresponding to 2-3 brains from each genotype was finely minced and resuspended in buffer A (25 mM HEPES, 1mg/ml glucose in 1X-HBSS) containing 1 Unit/ml papain and 10 µg/ml bovine pancreatic DNase. Following incubation for 30 minutes at 37°C in a shaking water bath at 280 rpm for tissue digestion, the tissue was extensively washed with ice-cold buffer A and forced through a 75 μm pore size sterile nylon. The resulting single-cell suspension was then subjected to centrifugation in a self-generated isotonic Percoll gradient at 14,160 rpm for 15 minutes at 4°C in a fixed-angle rotor (JLA 16.250). A lower Percoll gradient layer containing oligodendrocytes was collected and washed with ice-cold buffer A to remove residual Percoll. The cell pellets were then resuspended in serum-free chemically defined medium (N2 medium in DMEM/Ham’s F12) and subjected to differential attachment in a 100 mm tissue culture-treated dish for 20 minutes at 37°C to eliminate microglia and potential residual astrocytes. After gently swirling, non-attached and floating oligodendrocytes were pelleted down by centrifugation at 1000 rpm for 5 min at 4°C. and either resuspended in protein lysis buffer for protein extraction or Trizol for total RNA extraction.

##### Optic nerve collection

Optic nerves were also collected for RNA and protein extraction after the removal of the cerebral hemispheres.

#### 2.2.3. *Ormdl3* cKO verification and validation

Littermates were weaned, and tails were snipped for genotyping. Total DNA was extracted by adding 100µl of 50 mM sodium hydroxide to each tail sample. The samples were incubated at 95°C for 1 hour, with the tubes flicking every 20 min to ensure the tail was completely dissolved. Samples were cooled to room temperature, and 100 µl of 0.5 M sodium acetate was added to each tube to neutralize the sodium hydroxide, followed by immediate mixing. Samples were stored at -20°C until further use. The presence of *Cnp*-Cre was verified by PCR using primers specifically designed to amplify *Cnp*-Cre (primer details in **Supplementary Table 1)**. The successful *Ormdl3* cKO was validated by real-time PCR in isolated OLGs from P16 mice (primer details in **Supplementary Table 2).** The *Ormdl3* cKO in the cerebral hemispheres was re-verified by PCR. Here, total DNA was extracted from tissue, and the region around the loxP sites was amplified using specific primers.

#### 2.2.4. RNA isolation

Total RNA was extracted from isolated OLGs, optic nerves, and total cerebral hemispheres. The samples were resuspended in Trizol reagent and transferred to a pre-spun 2 ml heavy phase lock gel tube. To each tube, 200 µl chloroform/isoamyl alcohol (49:1) was added, followed by vigorous vortexing for 30 seconds. The samples were incubated at room temperature for 3 minutes, and phases were separated by centrifuging at 13,000 rpm for 15 minutes at 4°C. The upper clear phase was collected into a 1.5 ml microfuge tube, and 500 µl of isopropanol, 150 µl of 3M sodium acetate, and 2 µl of Glyco blue precipitate were added, followed by a quick, gentle vortex. The samples were incubated at -20°C for 20 minutes. RNA was pelleted by centrifuging at 13,000 rpm for 20 min at 4°C. RNA pellets were washed 3 times with 70% ice-cold ethanol by centrifugation at 13,000 rpm for 5 minutes. After the last wash, ethanol was carefully aspirated, and RNA pellets were dried under sterile conditions for 15 minutes. Dried pellets were resuspended using pre-warmed nuclease-free water (15-30 µl depending on pellet size). The RNA concentration was quantified using a Nanodrop 2000 (Thermo Scientific).

#### 2.2.5. cDNA preparation and qRT-PCR

cDNA was prepared using a high-capacity reverse transcriptase kit per the manufacturer’s protocol (Applied Biosystems). For each sample, 1 µg of total RNA was reverse-transcribed into single-stranded cDNA. Pre-designed FAM-labeled quantitative-PCR primer-probe sets and SYBR green dye-compatible pre-designed primers were used for the gene of interest. All the primers and primer-probe sets were ordered from IDT. For real-time PCR analysis, 5 -15 ng of cDNA per sample per gene was used along with PrimeTime Gene Expression Master Mix or Bio-Rad SYBR Green qPCR master mix. The CFX Connect real-time PCR detection system (Bio-Rad) was used to amplify the cDNA. Gene expression was calculated by the ΔΔCt method [40], and results were normalized to housekeeping genes and set relative to control gene expression levels.

#### 2.2.6. Myelin Isolation

Postnatal day 30 mice were used for the myelin isolation by density gradient centrifugation as previously described[41, 42]. Mice were anesthetized by using isoflurane, followed by the rapid decapitation. The cerebral hemispheres were collected and homogenized in 4.5 ml of 0.3M sucrose prepared in buffer A supplemented with 1X EDTA-free protease inhibitor cocktail. An aliquot of 100 µl of the total brain homogenate was set aside for protein quantification. The ultra-clear Beckman tubes were kept on ice, and 4.5 ml of 0.83 M sucrose in buffer A was added. The brain homogenates were carefully overlaid over 0.83 M sucrose. The samples were centrifuged at 75,000 g for 35 minutes at 4°C using a Beckman ultracentrifuge (20,000 RPM in SW41 Ti rotor). Carefully, tubes were removed from the centrifuge without disturbing the gradient. The interface was collected and re-homogenized in 15 ml of buffer A (20 mM Tris-HCl, pH 7.4 (hypotonic buffer A) + 5 mM EDTA), followed by centrifugation at 12,000 g for 15 minutes at 4°C (Avanti JLA; 9,000 RPM in JLA-16.250 rotor). The pellet was collected and resuspended in 0.3 M sucrose. The gradient centrifugation step was repeated by overlaying the resuspended pellet in 0.3 M sucrose over 0.83 M sucrose, followed by centrifugation at 75,000 g for 35 minutes at 4°C. The interface was collected in a tube, and 8 mL buffer A was added, followed by vigorous mixing and vortexing. The samples were centrifuged at 12,000 g for 15 minutes at 4°C. The final purified myelin pellet was resuspended in buffer A, containing 1X EDTA-free protease inhibitor cocktail, and aliquoted for future experiments. The successful isolation of pure myelin was verified through immunoblotting, which detected the myelin-specific proteins.

#### 2.2.7. Immunoblotting

Total cerebral hemisphere protein lysates and optic nerve protein lysates were prepared using 0.3M sucrose (prepared in 20 mM Tris-HCl, pH 7.4), and protein concentrations were quantified using Bradford reagent. Immunoblot samples were prepared by adding 5X Laemmli buffer to protein lysates, then incubating at 60°C for 30 minutes. An equal number of samples were electrophoresed on in-house prepared Tris-SDS gradient gels (a gradient of 10%, 12% and 15%). The separated proteins from the gel were transferred onto a pre-activated 0.22 μM PVDF membrane using the cold-wet transfer method. Blots were cut into two parts to probe for housekeeping protein and protein of interest. Blots were incubated with a 5% blotting-grade blocker for 1 hour at room temperature, then overnight at 4 °C with gentle agitation in primary antibodies. The next day, blots were washed 3 times with 1X-TBST for 10 minutes each, followed by overnight incubation with HRP-conjugated secondary antibodies. The next day, blots were washed 3 times with 1x-TBST for 10 minutes each, then with Milli-Q water for 15 minutes. The blots were visualized using an ECL-plus reagent per the manufacturer’s instructions and imaged on an Azure imager by Azure Biosystems.

#### 2.2.8. Electron Microscopy

On postnatal day 30, mice were transcardially perfused with 0.1 M Millonig’s buffer containing 5% glutaraldehyde and 4% paraformaldehyde, followed by whole-body post-fixation for 2 weeks at 4 degrees C. After two weeks, the brain and optic nerves were harvested, cut into longitudinal and cross-sectional sections, and incubated overnight in 0.1 M sodium cacodylate buffer. The next day, tissues were rinsed 3 times with cacodylate buffer for 10 minutes, followed by a 1-hour post-fixation in 1% osmium tetroxide (prepared in 0.1 M sodium cacodylate buffer) with gentle shaking at room temperature. Next, tissues were rinsed 3 times with 0.1 M sodium cacodylate buffer (5 minutes per wash), followed by slow dehydration using ethanol at different concentrations. The ethanol concentrations used were 30%, 50%, 70%, 90%, and 95% (2 rinses, 5 minutes each), one rinse with 100% ethanol for 5 minutes, and three rinses with 100% ethanol for 10 minutes each. Finally, tissues were rinsed twice with propylene oxide for 30 minutes at room temperature. The tissues were incubated in a solution of 1 part propylene oxide and 1 part PolyBed resin overnight at room temperature with continuous agitation. The next day, tissues were incubated in 100% PolyBed resin for 24 hours with continuous mixing. Finally, tissues were embedded in Poly/Bed 812 (Poly/Bed + dodecenylsuccinic anhydride, nadic methyl anhydride, DMP-30) by polymerizing in an oven at 60°C for 36 hours. The tissues were sectioned at 70 nm and stained with lead citrate and uranyl acetate. The sections were imaged using a JEOL JEM1400 PLUS transmission electron microscope at the VCU Microscopy Core Facility.

#### 2.2.9. Analysis of Transmission Electron Microscopy images

Transversely sectioned tissues were used to measure g-ratios, myelin thickness, and axon diameters using ImageJ. At least 10 electron micrograph images were captured at 5000X per mouse. For all calculations, we focused on axons without tissue fixation artifacts. EM image quantification was done blindly.

#### 2.2.10. Immunohistochemistry and RNAscope

Immunohistochemistry and RNAscope were performed as outlined by Spencer et al., 2022[43], and tissue sections were prepared as described previously[44, 45]. Briefly, mice were anesthetized by intraperitoneal injection of 0.8ml/20g (of mouse body weight) 2-2-2 tribromoethanol and then transcardially perfused with 4% paraformaldehyde in 0.1 M Millonig’s phosphate buffer[46]. Brains were removed, post-fixed for 24 hours in perfusion fixative, cryoprotected by immersion in 30% sucrose in PBS for 48 hours, and then embedded and frozen in Tissue-Tek O.C.T. compound.

For immunohistochemistry, serial coronal (P16) or sagittal (P28) sections (40 μm) were prepared using a Leica CM1850 cryostat (Leica Biosystems, Buffalo Grove, IL) and stored at −80°C. Tissue sections were permeabilized for 10 minutes on ice in ice-cold acetone and blocked for 15 min at room temperature in PBS containing 10% Triton X-100 and 10% normal goat serum. Primary antibodies were diluted in PBS containing 0.2% Triton X-100 and 10% normal goat serum, and sections were incubated for 24 hours at room temperature, followed by incubation with secondary antibodies for 90 min at room temperature. Nuclei were counterstained using Hoechst 33342, and sections were mounted using Vectashield. The following antibodies were used: anti-Olig2, CC1, goat anti-Chicken IgY, Alexa Fluor 633, and goat anti-Mouse IgG2b, Alexa Fluor 568.

For RNAscope, serial sagittal sections (15-μm) were prepared using a Leica CM 1850 cryostat (Leica Biosystems, Buffalo Grove, IL), sections were mounted on Leica Bond Plus slides (Cat. No. S21.2113.A, Leica Biosystems, Deer Park, IL) and post-fixed as follows: 60°C for 30 min, 4% paraformaldehyde in PBS for 15 min at 4°C, 50% ethanol for 5 min at room temperature, 70% ethanol for 5 min at room temperature, 100% ethanol for 5 min at room temperature (twice), air dried for 5 min, and stored at −80°C. RNAScope was performed using a Leica Biosystems Bond RX automated immunohistochemistry*/in situ* hybridization staining system (Leica Biosystems, Deer Park, IL located within VCU’s Tissue and Data Acquisition and Analysis Core and the following RNAscope 2.5 LS Probe probes (all from Advanced Cell Diagnostics, Inc., Newark, CA): Mm-*Olig2*-C2, Mm-*Plp1*-C4, Mm-*Mog*-C3, for fluorescent detection, RNAscope LS Fluorescent Reagent and RNAscope LS 4-Plex Ancilliary kits were used in combination with the following Opal fluorophores (Akoya Biosciences, Marlborough, MA): Opal 690, Opal 620, Opal 520. Images were obtained using a Vectra PhenoImager HT imaging system (Akoya Biosciences, Marlborough, MA) located within VCU’s Tissue and Data Acquisition and Analysis Core. For quantification, double-positive cells were manually counted using the Fiji/ImageJ image-processing package and its Cell Counter plugin [47].

#### 2.2.11. Lipidomics

##### 2.2.11.1. Extraction of Sphingolipids

Tissue homogenates or cell pellets were collected in 13 x 100 mm borosilicate tubes with a Teflon-lined cap. Then, 2 mL of CH3OH was added along with the internal standard cocktail (250 pmol of each species dissolved in a final total volume of 10 mL of ethanol:methanol:water (7:2:1). The mixture was dispersed using an ultrasonicator at room temperature for 30 s. Then 1 mL of CHCl3 was added, and the test tubes were recapped. This single-phase mixture was incubated at 48 °C overnight. The extract was centrifuged using a tabletop centrifuge, and the supernatant was removed using a Pasteur pipette and transferred to a new tube. The extract was reduced to dryness using a Speed Vac. The dried residue was reconstituted in 0.5 ml of the starting mobile-phase solvent for LC-MS/MS analysis, sonicated for ca 15 sec, then centrifuged for 5 min in a tabletop centrifuge before transferring the clear supernatant to the autoinjector vial for analysis.

##### 2.2.11.2. LC-MS/MS of sphingoid bases, sphingoid base 1-phosphates, and complex sphingolipids

These compounds were separated by reverse-phase LC using a Supelco 2.1 (i.d.) x 50 mm Ascentis Express C18 column (Sigma, St. Louis, MO) and a binary solvent system at a flow rate of 0.5 mL/min with a column oven set to 35°C. Before injection of the sample, the column was equilibrated for 0.5 min with a solvent mixture of 95% Moble phase A1 (CH3OH/H2O/HCOOH, 58/41/1, v/v/v, with 5 mM ammonium formate) and 5% Mobile phase B1 (CH3OH/HCOOH, 99/1, v/v, with 5 mM ammonium formate). After sample injection (typically 40 μL), the A1/B1 ratio was maintained at 95/5 for 2.25 min, followed by a linear gradient to 100% B1 over 1.5 min, which was held at 100% B1 for 5.5 min, followed by a 0.5 min gradient return to 95/5 A1/B1. The column was re-equilibrated with 95:5 A1/B1 for 0.5 min before the next run.

##### 2.2.11.3. LC-MS/MS of Glucosylceramide and Galactosylceramide

Because glucosylceramide and galactosylceramide co-elute using the above method, biological samples containing both can be analyzed by a separate method. Dried samples are re-dissolved in CH3CN/CH3OH/H3CCOOH (97:2:1) (v,v,v) with 5 mM ammonium acetate. The LC-Si column (Supelco 2.1 x 250 mm LC-Si) was pre-equilibrated with CH3CN/CH3OH/H3CCOOH (97:2:1) (v/v/v) with 5 mM ammonium acetate for 1.0 min at 1.5 mL per min. The sample was injected, and the column was isocratically eluted for 8 min. GlcCer elutes at 2.56 min and GalCer at 3.12 min using this isocratic normal-phase system; however, column age and previous sample load can influence the retention times of HexCers by this method. Periodic confirmation of retention time using internal standards enables monitoring of column stability and subsequent effectiveness.

##### 2.2.11.4. LC-MS/MS of Sulfatide

The sulfatide internal standard (d18:1/C12:0 sulfatide), Avanti Polar Lipids (Alabaster, AL). In experiments where sulfatide analysis was required, 250 pmol was added with the other internal standard during extraction. Sulfatides were separated by reverse-phase LC using a Supelco 2.1 (i.d.) x 50 mm Ascentis Express C18 column (Sigma, St. Louis, MO) and a binary solvent system at a flow rate of 0.7 mL/min with a column oven set to 60°C. Before injection of the sample, the column was equilibrated for 0.5 min with a solvent mixture of 99% Moble phase A1 (CH3OH/H2O/HCOOH, 65/34/1, v/v/v) and 1% Mobile phase B1 (CH3OH/HCOOH, 99/1, v/v). After sample injection (typically 10 μL), the A1/B1 ratio was maintained at 99/1 for 3.0 min, followed by a linear gradient to 100% B1 over 2.25 min, which was held at 100% B1 for 4.5 min, followed by a 0.5 min gradient return to 99/1 A1/B1. The column was re-equilibrated with 99:1 A1/B1 for 0.5 min before the next run. Sulfatides were analyzed in negative ion mode, using an m/z 240.9 product ion (indicative of the sulfated galactose) for quantitation. Previously, sulfatide standards were used to confirm LC retention in addition to N-acyl fatty acid product ions.

#### 2.2.12. Cholesterol Assay

The cholesterol assay was performed using the Amplex Red Cholesterol Assay Kit (Invitrogen #12216). The assay was performed following the manufacturer’s protocol. In brief, the assay was performed on a 96-well black plate (ThermoFisher #14-245-197A). The cholesterol standard curve was prepared by diluting the 2mg/ml reference standard with 1X Reaction buffer to generate concentrations ranging from 0 to 8 µg/ml. As a negative control, a reaction buffer without a reference standard was used. For the assay, 0.5 µg of total myelin samples was diluted in 1X Reaction buffer. 50 µl of each sample and reference standard was pipetted in triplicate into a 96-well plate. A working solution of Amplex Red reagent was prepared as follows: 2 U/mL HRP and 2 U/mL cholesterol oxidase, obtained by adding 75 μl of Amplex® Red reagent stock solution. 50 µl of Amplex Red reagent stock solution was added to each well, and the plate was incubated at 37°C for 30 minutes. After 30 minutes, fluorescence was measured using a microplate reader with excitation at 560 nm and emission at 590 nm. The background was subtracted from the samples by subtracting the values of the standard and the samples from those of the no-cholesterol controls.

#### 2.2.13. Preparation of Rat Oligodendrocyte Cell Cultures and siRNA knockdown of *Ormdl3*

Sprague-Dawley female rats and their natural 7-day-old pups (10 pups per litter) were obtained from Charles River Laboratories (Wilmington, MA). Animals were housed for 24 hours under a light/dark cycle and temperature-controlled conditions, with access to food and water. Studies were conducted in accordance with the National Institutes of Health *Guide for the Care and Use of Laboratory Animals* and under protocols approved by the Animal Care and Use Committee of Virginia Commonwealth University, Richmond, VA.

OLGs were isolated from 8-day-old pups using a Percoll gradient centrifugation and differential cell attachment protocol as previously described[39] and detailed above under 2.2.2. Isolated oligodendrocytes were seeded on 48-well Matrigel pre-coated plates and maintained in 250 µl chemically defined medium [DMEM/F12 (1:1) medium with high glucose and L-glutamine, pH 7.4, containing N2 supplement (Invitrogen #17502001), 1 mg/ml endotoxin-free fatty acid-free bovine serum albumin, and 30 nM triiodothyronine (T3)]. siRNA knockdown experiments were carried out on the following day five hours after a fresh medium change.

In brief, 20 pmol of each of the three *Ormdl3* siRNAs (Integrated DNA Technologies, # Rn.PT.58.33915101, Rn.PT.58.9412021, Rn.PT.58.5987952.gs) and, as a control, 60 pmol of scrambled siRNA (Integrated DNA Technologies #51-01-19-09) were diluted with 150 µl Opti-Mem. Next, RNAiMax (Invitrogen #13778100) was diluted with Opti-Mem and incubated for 5 min at room temperature. After 5 min, diluted siRNAs were added to diluted RNAiMax, and the mixture was incubated for 15 min at room temperature. After 15 min, the transfection mix was filter sterilized through 0.22 µm syringe filters (Millipore #WHA9913-2502). A 25 µl transfection mix was added to each well. After every 24 hours, 150 µl of the medium was replaced with fresh medium. For RNA isolation and real-time experiments, cells were collected after 4 days of transfection, and for protein extraction, after 5 days of transfection. Real-time PCR and immunoblotting were performed as described in previous sections.

## 3. Results

### 3.1. Validation of Oligodendrocyte-Specific *Ormdl3* Conditional Knockout in Postnatal Day 16 Mice

The OLG-specific *Ormdl3* cKO mice were generated using the Cre-lox gene-editing system [48]. Mice expressing Cre recombinase under the control of the 2’,3’-cyclic nucleotide 3-phosphodiesterase (*CNPase*) promoter, an OLG-specific enzyme, were crossed with mice carrying LoxP sequences flanking exon 2 to exon 4 of *Ormdl3* (**Fig. 1B-1C**). *Cnp* is expressed during early brain development, with peak enzyme activity before postnatal day 10 [37]. As a result of crossbreeding, the expression of *Ormdl3* in OLGs present in the CNS (spinal cord, brain, and optic nerves) will be affected before postnatal day (P) 3, resulting in the generation of OLG-specific cKO mice. To verify the deletion of *Ormdl3*, we isolated OLGs from P16 mice. RT-qPCR was performed to measure *Ormdl3* mRNA levels using *Ormdl3* qPCR probes specific to the deleted region (exon 2 to exon 4). As expected, *Ormdl3* mRNA expression was not detected in *Ormdl3* cKO mice (OLG^ΔORMDL3^) compared to the WT (*Ormdl3^floxed/floxed^)* group (**Fig. 1D**). To confirm that *Ormdl3* cKO is specific to OLGs, we also collected microglia from the same mice and measured *Ormdl3* mRNA levels. We did not observe any changes in *Ormdl3* mRNA levels in microglia isolated from *Ormdl3* cKO mice compared to WT mice. Furthermore, PCR amplification of *Ormdl3* from genomic DNA isolated from the total cerebral hemispheres of *Ormdl3* cKO mice confirmed that Cre-lox recombination occurs in the P16 mouse CNS (**Fig. 1E**). The wild-type full-length *Ormdl3* band in cKO samples was detectable at a low level compared with the WT sample (**Sup Fig. 1A**). The abundance of ORMDL protein in OLGs is very low, which precludes us from detecting protein levels even in WT mice.

### 3.2. *Ormdl3* Deletion Elevates Myelin Sphingolipids in Postnatal Day 35 Mice

To investigate the effect of *Ormdl3* deletion in OLGs on myelin SLs, we isolated myelin from the total brain homogenates (TBH) of postnatal day 30 (P30) and day 35 (P35) mice from the WT and *Ormdl3* cKO groups of both sexes. We verified the purity of isolated myelin by immunoblotting (**Figs. 2A-2B**). Myelin proteins, such as MBP and MOG, were found to be enriched in the myelin but not the astrocyte marker Glial Fibrillary Acidic Protein (GFAP) [49] in both WT and cKO mouse groups. We measured the steady-state levels of canonical SLs, including sulfatides (non-hydroxylated and hydroxylated), sphingomyelin, galactosylceramides, ceramides, sphingoid long-chain bases (sphingosine, sphingosine-1-phosphate, and dihydrosphingosine), monohexosylceramides (sum of glucosylceramides and galactosylceramides), and glucosylceramides in both WT and *Ormdl3* cKO mice (both sexes). We analyzed species of different chain lengths and will refer to total levels as the sum of acyl chain lengths (C14:0, C16:0, C18:1, C18:0, C20:0, C22:0, C24:1, C24:0, C26:1, and C26:0). To enhance clarity, the results for sulfatide levels are compiled in **Fig. 2H (Table 1B)**.

**Figure 2:**
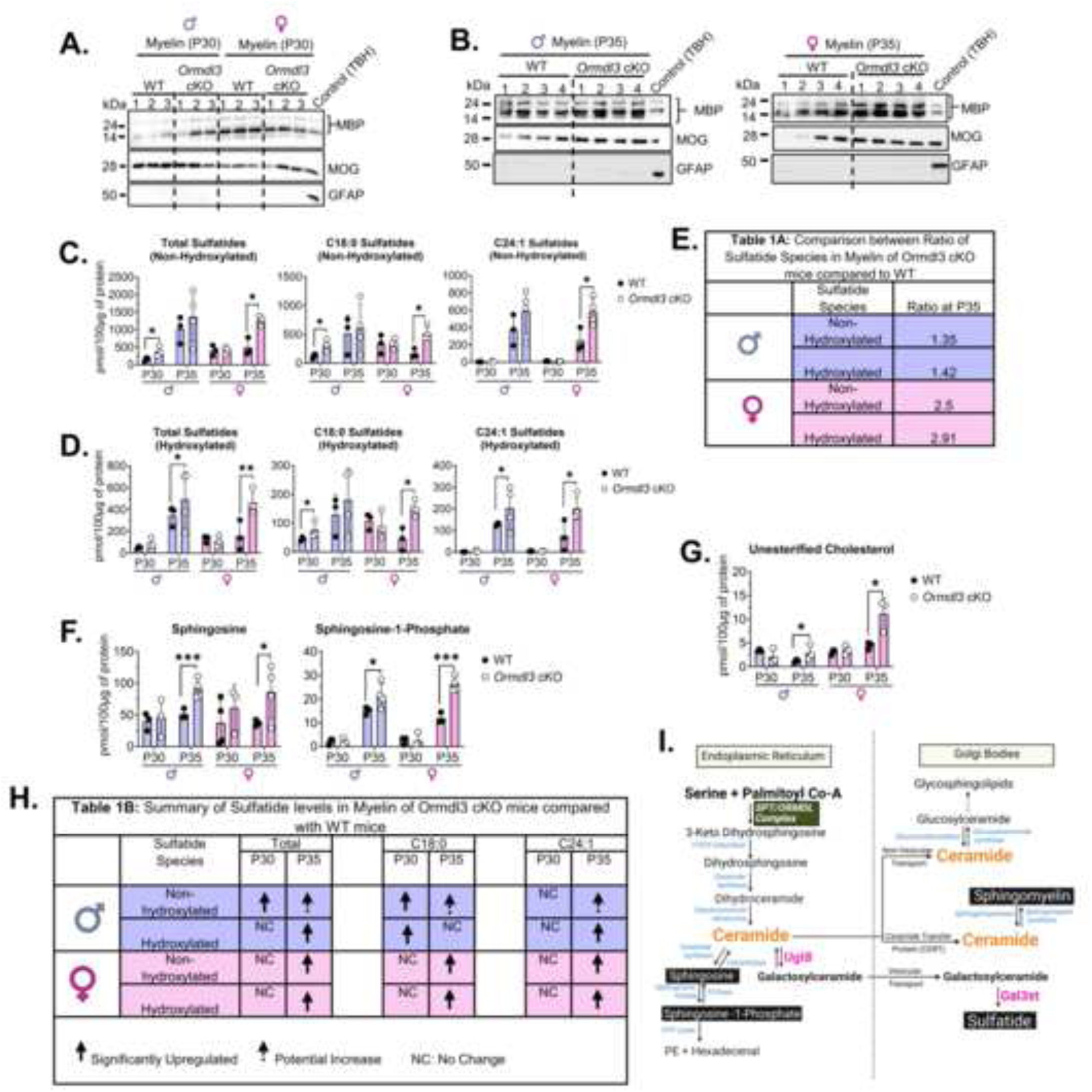
Altered myelin sulfatide, sphingoid bases, and cholesterol levels in oligodendrocyte-specific *Ormdl3* cKO mice around postnatal day 35. Total myelin was isolated from the brains (TBH) of mice on postnatal days 30 (P30) and 35 (P35) from WT and *Ormdl3* cKO groups. Purity of the isolated myelin was checked by immunoblotting **(A-B)** at P30 and P35 in both sexes of mice. In brief, 0.2 µg of myelin sheath was loaded into each lane of in-house-made gradient Tris-SDS polyacrylamide gels as described in the methods section. Myelin Basic Protein (MBP) and Myelin Oligodendrocyte Glycoprotein (MOG) antibodies were used to confirm the purity of myelin. Glial Fibrillary Acidic Protein (GFAP) was used as a negative control, and TBH was used as a positive control. The steady-state levels of **C.)** total non-hydroxylated sulfatides, and C18:0 and C24:1 acyl chain length non-hydroxylated sulfatide species, **D.)** total hydroxylated sulfatides, and C18:0 and C24:1 acyl chain length hydroxylated sulfatide species in the myelin of WT and cKO mice at P30 and P35, in both sexes, respectively. **E.) Table 1A:** The ratio of hydroxylated vs non-hydroxylated sulfatide species at P35 in both WT and *Ormdl3* cKO mice, in both sexes, respectively. The ratio was calculated by dividing the pmol of sulfatides (non-hydroxylated and hydroxylated) per 100 μg of protein from ORMDL3 cKO mice by the pmol of sulfatides (both non-hydroxylated and hydroxylated) per 100 μg of protein from WT mice. **F.)** The steady-state levels of sphingosine and sphingosine-1-phosphate in WT and *Ormdl3* cKO mice, at P30 and P35, in both sexes, respectively. **G.)** The levels of unesterified cholesterol in the myelin of WT and *Ormdl3* cKO mice at P30 and P35, in both sexes, are shown. **H.) Table 1B:** Summary of the results from panels **C** and **D**. **I.)** Cartoon representation of *the de novo* Sphingolipid biosynthesis pathway, highlighting the Sphingolipid species (black boxes) that are increased in *Ormdl3* cKO mice compared with WT groups. The sphingolipid levels were measured by Mass Spectrometry as described in the methods section. The lipidomic data are graphed as picomoles of lipids per 100 µg of protein. Shown are the mean ± SD, n = 3-4. Statistical significance was tested by the Student’s two-tailed t-test. *Asterisks* denote significance, p <0.05 = *, p <0.005 = ** and p <0.0005 = ***. Purple-colored bars denote males and pink-colored bars denote females.

In general, there is a clear increase in both hydroxylated and non-hydroxylated sulfatides on day 35 due to ORMDL3 deletion, with some sex differences. At P30, the results are less consistent. At P30, the levels of total non-hydroxylated sulfatides were significantly upregulated in cKO male mice, but not in cKO female mice, compared to WT groups (**Fig. 2C**). By P35, male cKO mice showed a trend toward increased total non-hydroxylated sulfatides, whereas in female cKO mice, these levels were significantly elevated compared with WT groups. Our analysis of different chain-length species revealed that the predominant species at P30 is C18:0, whereas at P35, C18:0 and C24:1 are predominant in both sexes of WT and cKO mice (**Fig. 2C and Sup Figs. 1A-1B**). At P30, the C18:0 species is significantly upregulated in male cKO mice, but no changes were observed in female cKO mice compared to WT groups. By P35, both chain-length species (C18:0 and C24:1) are significantly upregulated in cKO female mice, and trends toward increased levels are observed in cKO male mice compared to WT.

The total hydroxylated sulfatides are also significantly upregulated in both sexes of *Ormdl3* cKO mice compared to WT at P35 but not at P30 (**Fig. 2D**). Analysis of different chain lengths of hydroxylated sulfatides revealed that C18:0 is the predominant species at P30, and by P35, the chain length shifts to C18:0 and C24:1 in both WT and cKO mice of both sexes (**Fig. 2D and Sup Figs. 1C-1D**). Both chain lengths (C18:0 and C24:1) are significantly upregulated in cKO mice of both sexes compared to WT groups. The ratio of non-hydroxylated vs hydroxylated is not altered in cKO mice (both sexes) compared to WT groups, suggesting that sulfatide species are proportionally increased in cKO mice **Fig. 2E**; **Table 1A)**. The steady-state levels of sphingoid long-chain bases (sphingosine and sphingosine-1-phosphate) are significantly upregulated at P35 in cKO mice (both sexes) but not at P30 compared with WT groups (**Fig. 2F**). However, no changes were observed in dihydrosphingosine levels at both P30 and P35 in cKO mice of both sexes compared to WT (**Sup Fig. 3A**). Upon measuring unesterified cholesterol levels, we observed that at P35, levels were significantly upregulated in cKO mice (both sexes), but not at P30 (**Fig. 2G**). Cholesterol is the most abundant lipid in the myelin sheath, which is essential for the growth and maintenance of myelin [50].

Effects on sphingomyelin levels were less clear. While total levels of sphingomyelin in myelin were either not or only slightly significantly different between WT and cKO mice, there were some differences in male mice of specific molecular species at day 35. At P30, the predominant chain-length species are C18:0 and C24:1, and by P35, preference shifts to C18:0 and C20:0 in both WT and cKO mice of both sexes (**Sup Figs. 4B-4C**). At P35, both C18:0 and C24:1 species are significantly upregulated in male cKO mice compared to WT (**Sup Fig. 4B**).

We did not observe any effect of *Ormdl3* cKO on myelin total ceramides (**Sup Fig. 5A**), although there was a striking increase in the ceramide content of myelin in both WT and cKO animals between P30 and P35 due to a dramatic increase in C24:1 levels. Total galactosylceramides (**Sup Fig. 6A**) and total glucosylceramide levels (Sup Fig. 7A) at either age (P30 and P35) in both sexes were unchanged in the cKO animals compared with WT. The predominant ceramide and galactosylceramide species at P30 are C24:1, and by P35, C24:1 remains the predominant species in both male and female WT and cKO mice (**Sup Figs. 5B-5C and Sup Figs. 6B-6C**). The predominant species of glucosylceramides at P30 are C24:1 and C24:0; by P35, the preference shifts to only C24:1 in both male and female WT and cKO mice (**Sup Figs. 7B-7C**). The above results suggest that during myelin development, the incorporation of SL species with varying chain lengths changes dramatically in mice from P30 to P35, with the incorporation of long-chain bases into the myelin SLs. Similarly, the *Ormdl3* cKO mice exhibited a comparable pattern of changes in the incorporation of species with differing chain lengths. Notably, there was a selective increase in the levels of sulfatide, sphingosine, sphingosine-1-phosphate, and, to some extent, sphingomyelin in the cKO mice compared to the WT groups (**Fig. 2H and 2I**).

### 3.3. *Ormdl3* cKO Mice Optic Nerve Shows a Coordinated Increase in Myelin Thickness and Larger-Caliber Axons on Postnatal Day 30

To investigate the ultrastructural effects of *Ormdl3* cKO on myelin in the CNS, we examined optic nerve cross-sections using electron microscopy. We chose optic nerves for these experiments because myelination is uniform along the nerve fiber, unlike in the brain, where different regions myelinate at different times during development [51]. Our analysis showed that cKO mice had increased myelin thickness (**Figs. 3A, 3B, and 3D**). This was accompanied by an increase in axon diameter (**Figs. 3A, 3C, and 3E**). However, the G-ratio (ratio of axon diameter to outer fiber diameter) (**Fig. 3F**) was not significantly different between WT and cKO *mice*. The g-ratio is a widely utilized parameter in morphological studies to evaluate the optimization of axonal myelination and to assess the relative thickness of the myelin sheath [52]. We also compared the electron microscopy results between male and female mice and observed no differences, indicating no sexual dimorphism.

**Figure 3:**
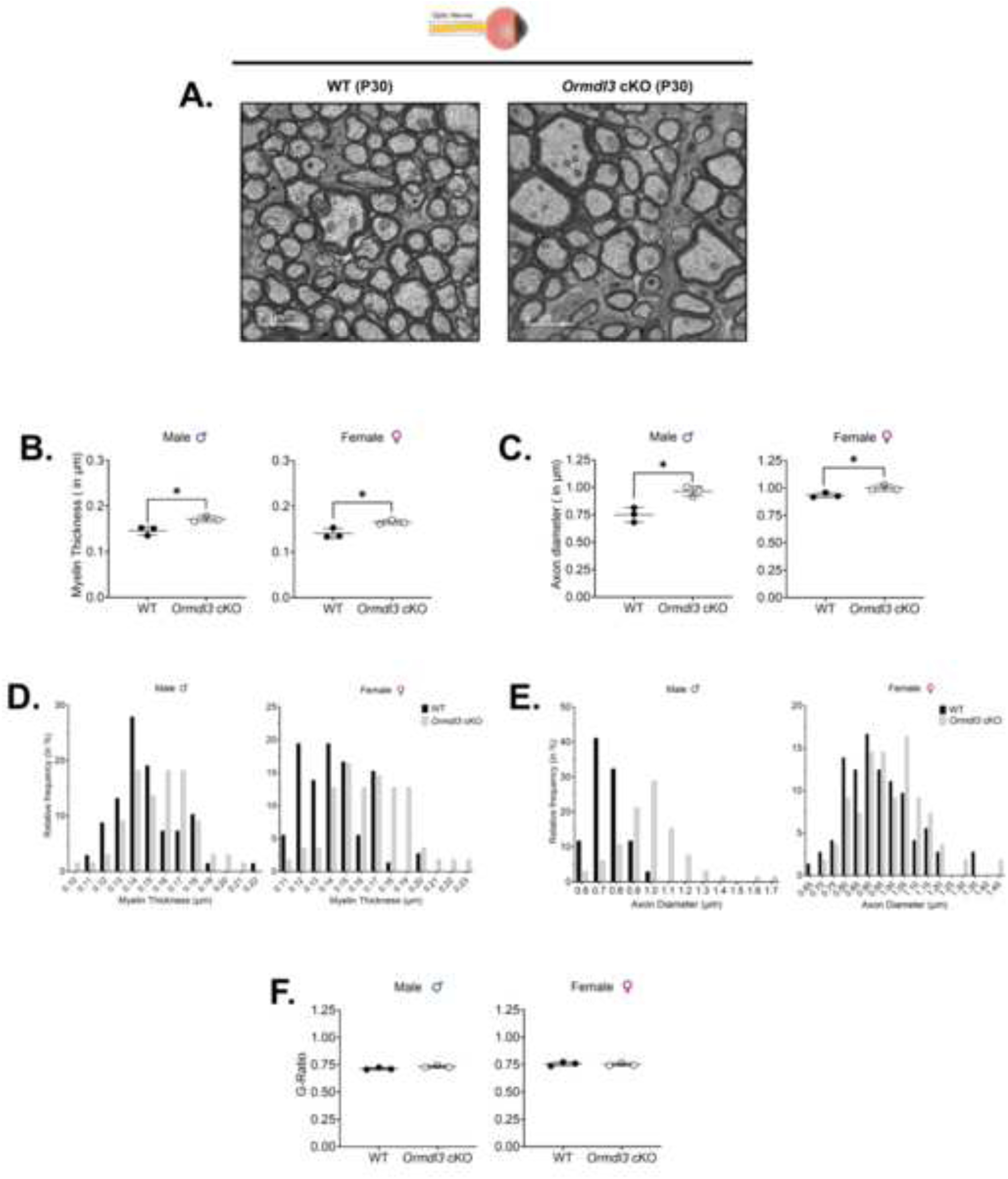
Effect of *Ormdl3* deletion on the myelin’s ultrastructure in optic nerves. **A.)** Representative electron microscopy image of the optic nerve’s cross-sectional area (female). The scale bar was set to 2µm. **B.)** The quantification of the myelin thickness in WT and *Ormdl3* cKO mice of both genders. Myelin thickness was measured by dividing the circumference of the axon without myelin by the circumference of the axon with myelin. **C.)** The quantification of the axon caliber in WT and *Ormdl3* cKO mice of both genders. Axon Caliber was measured by measuring the circumference of the axon without myelin. **D.) and E.)** Frequency distribution curve of myelin thickness in WT and *Ormdl3* cKO mice. **E.)** Frequency distribution curve of axon diameter in WT and *Ormdl3* cKO mice. **D.)** Analysis of G-ratio in WT and *Ormdl3* cKO. The G-ratio is calculated by dividing the radius of the axon without myelin (r) by the radius of the axon with myelin (R). Shown are the mean ± SD, WT (3 males and 3 females), and *Ormdl3* cKO mice (3 males and 3 females). From each mouse, 10 images were captured, and from each image, at least 12-25 axons were analyzed. For panels **B**, **C**, and **E**, myelin thickness, axon diameter, and g-ratio were quantified from 10 images, and the quantification of 10 images was then averaged to obtain one data point per mouse. For the frequency distribution curves of myelin thickness and axon diameter, binning was performed using GraphPad Prism. Percentages were calculated based on at least 100 axons per mouse. Data from three WTs and three *Ormdl3* cKO mice were grouped to generate the final graph. Statistical significance was tested by the Student’s two-tailed *t*-test. *Asterisks* denote significance, *p* <0.05 = *.

### 3.4. Myelin Protein Expression in *Ormdl3* cKO Mice is Sex- and Region-Dependent

Lipidomic analysis of myelin SL in the brain at P35 and electron microscopy studies of optic nerves at P30 indicate an increase in the production of SL in *Ormdl3* cKO mice. Surprisingly, we also find increased levels of myelin protein in the total brain at P35 (**Fig. 4A**). At this time point, MBP and MOG protein levels were significantly increased in both male and female cKO mice compared to WT groups. PLP levels may be increased in the brain of male cKO mice, but the scatter in the data precludes a definitive assessment. No changes in PLP levels were observed in female cKO mice when compared to WT. These changes were not evident in the total brain at P30, in which protein levels were either unchanged or slightly decreased (MBP and PLP) only in female mice (**Sup Fig. 8A**). In the optic nerve at P30, we observed increased MBP levels in female cKO mice and, potentially, an elevation of PLP in both male and female mice. Interestingly, there is a slight decline in MOG levels in female cKO mice (**Fig. 4B**). We also measured myelin proteins at an early stage of myelin development (P16) in both the total brain and the optic nerve. No differences were observed in cKO mice compared with WT (**Sup Figs. 8B and 8C**).

**Figure 4:**
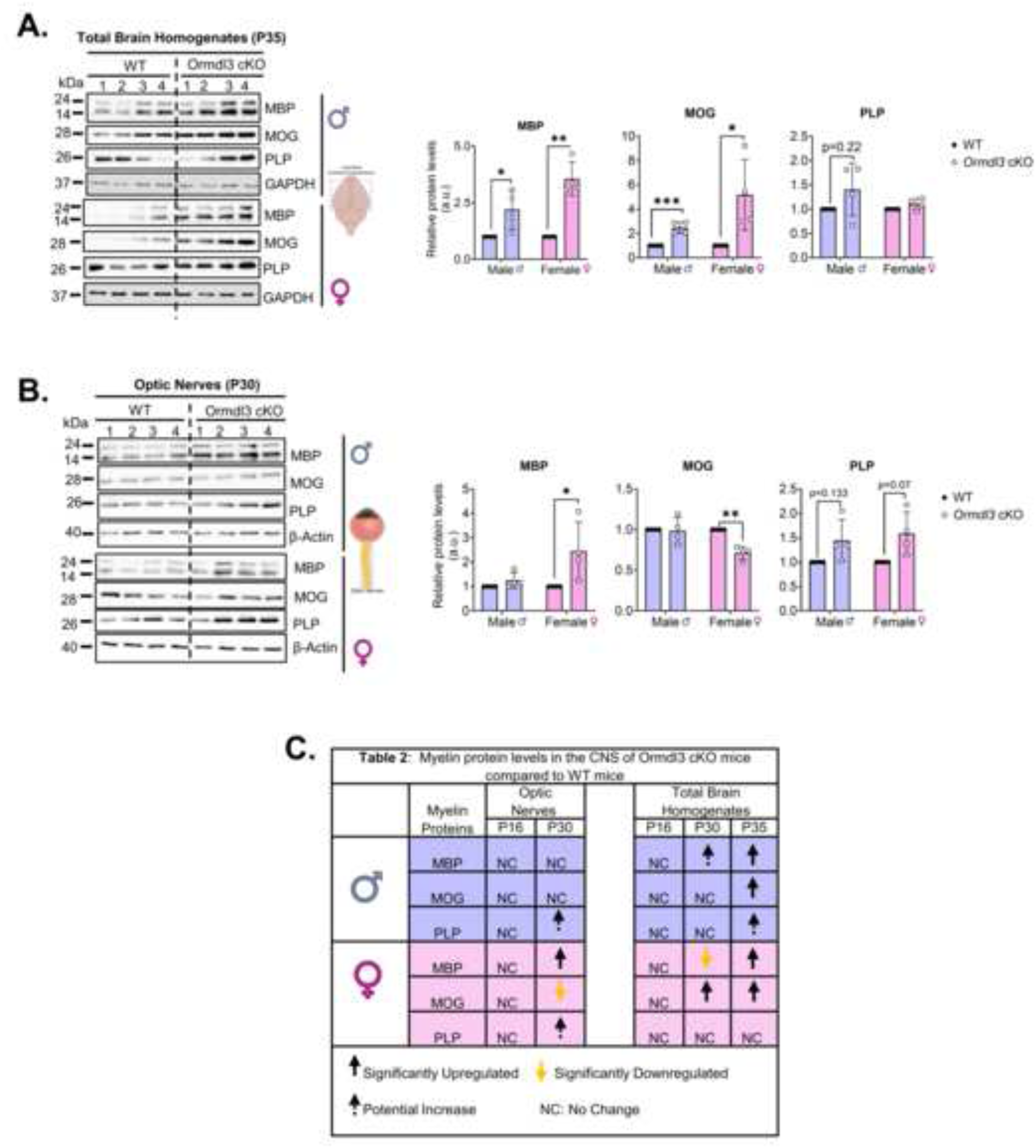
Effect of oligodendrocyte-specific *Ormdl3* deletion on myelin protein expression in the brain and optic nerve. Immunoblot analysis of myelin proteins: **A)** at P35 in total brain lysates with quantification of MBP, MOG, and PLP; **B)** at P30 in optic nerve lysates with quantification of MBP, MOG, and PLP, respectively. **C.) Table 2**: Summary of the results from panels **A** and **B**. In brief, 1-2 µg of lysate was loaded into each lane of in-house-made gradient Tris-SDS polyacrylamide gels as described in the methods section. The blots were then probed with MBP, MOG, PLP, GAPDH, and β-actin. GAPDH and β-actin were used as loading controls. The band intensity was then quantified by using Quantity One software. Data were normalized to GAPDH or β-actin and then set relative to WT. Data were shown as the mean ± SD; n=3–4. Statistical significance was tested by the Student’s two-tailed t-test. *Asterisks* denote significance, p <0.05 = *, p <0.005 = ** and p <0.0005 = ***. Here, we refer to the brain as the cerebral hemisphere, excluding the cerebellum.

### 3.5. Transcriptional Upregulation of Myelin Protein Genes in *Ormdl3* cKO Mice is Developmental Stage-Dependent

To assess whether the elevated levels of myelin proteins resulted from increased gene expression, we measured *Mbp, Mog,* and *Plp1* genes at various developmental stages in both the optic nerves and the brain. Results are from a mixed pool of male and female animals, as we did not observe any differences between the two genders. Additionally, we examined the mRNA levels of these myelin protein genes in brain-derived OLGs.

In the brain, mRNA levels were essentially identical between WT and cKO animals at all time points except at day 25, where we observed a striking increase in *Mbp* and *Mog* mRNA levels in cKO animals relative to WT animals (**Fig 5A**). There were no observed differences in *Plp1* expression. The *Mbp* and *Mog* expression data from total brain homogenates are supported by more limited data from oligodendrocytes isolated from the brain, which showed no differences between WT and cKO animals in *Mbp* or *Mog* expression at either day 16 or day 30 (**Fig 5B**). However, Plp1 expression decreased dramatically in cKO animals on day 16. This discrepancy in *Plp1* mRNA levels is addressed in the discussion.

**Figure 5:**
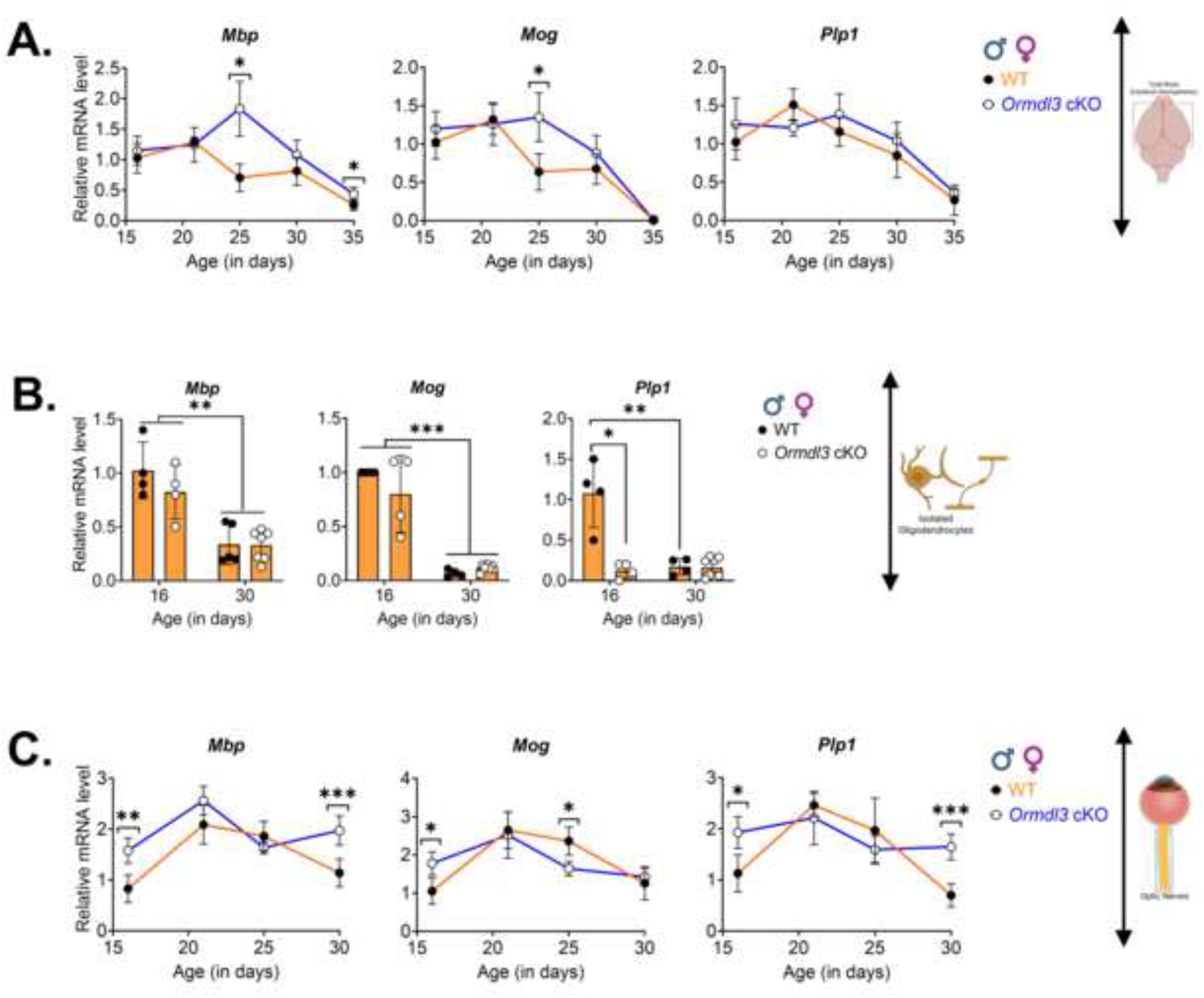
Effect of *Ormdl3* deletion in oligodendrocytes on myelin-associated protein mRNA levels. Shown are mRNA levels of **A.)** *Mbp*, *Mog,* and *Plp1* in the total brain isolated from WT and *Ormdl3* cKO mice at P16, P21, P25, P30, and P35 (mixed pool of sexes). Shown are the mRNA levels of **B.)** *Mbp*, *Mog,* and *Plp1* in oligodendrocytes isolated from WT and *Ormdl3* cKO mice at P16 and P30 (mixed pool of sexes). Shown are the mRNA levels of **C.)** *Mbp*, *Mog,* and *Plp1* in optic nerves isolated from WT and *Ormdl3* cKO mice at P16, P21, P25, and P30 (mixed pool of sexes). Total RNA was isolated as described in the methods section, and RT-qPCR was performed to measure mRNA levels. Data were normalized to either *Gapdh* or *Hprt* and set to WT (P16) as the reference. Data were presented as mean ± SD; n = 3–8 (a mixed pool of males and females). Statistical significance was tested by the Student’s two-tailed t-test. *Asterisks* denote significance, p <0.05 = *and p <0.0005 = ***. HPRT: Hypoxanthine phosphoribosyl transferase 1.

In the optic nerve, the picture was more complex. *Mbp* mRNA levels in the cKO animals were elevated relative to those of WT animals on days 16 and 30, but not in between (**Fig 5C**). *Mog* levels were slightly increased at days 16 and 25 in the cKO animals relative to WT. *Plp1* expression was similar to *Mbp* in that mRNA levels were increased in cKO vs WT animals at days 16 and 30.

### 3.6. *Ormdl3* deletion in Oligodendrocytes exhibits sex-specific modulation of Sulfatide and Cholesterol Metabolism

Our results indicate that the increase in SL production acts through an unknown mechanism to coordinately elevate levels of certain myelin proteins. To establish whether increased SPT activity, as a result of ORMDL3 deletion, has effects on other elements of the SL metabolic pathway, we measured the mRNA and protein levels of two key enzymes in the myelin SL biosynthesis pathway: UDP glycosyltransferase 8 (UGT8), which produces galactosylceramides, and galactose-3-O-sulfotransferase 1 (GAL3ST1), which uses galactosylceramides to synthesize sulfatides, which are enriched in myelin (**Fig. 6A**) [36]. When protein levels were measured in the total brain and optic nerve of P30 mice, we observed that ORMDL3 deletion induced an elevation of both UGT8 and GAL3ST1 levels in male, but not female, mice (**Figs. 6B and 6C**). These findings are summarized in **Fig. 6D** (**Table 4A**) for clarity. mRNA levels of *Gal3st1* were somewhat increased in cKO vs WT animals at day 30 in total brain but not isolated OLGs or optic nerve, but otherwise we did not observe any differences at day 30 between cKO and WT animals in the mRNA levels for *Ugt8* in either total brain, optic nerves, or isolated OLGs (**Figs. 6E and 6F**). These findings are summarized in **Fig. 6I** (**Table 4B)**

**Figure 6:**
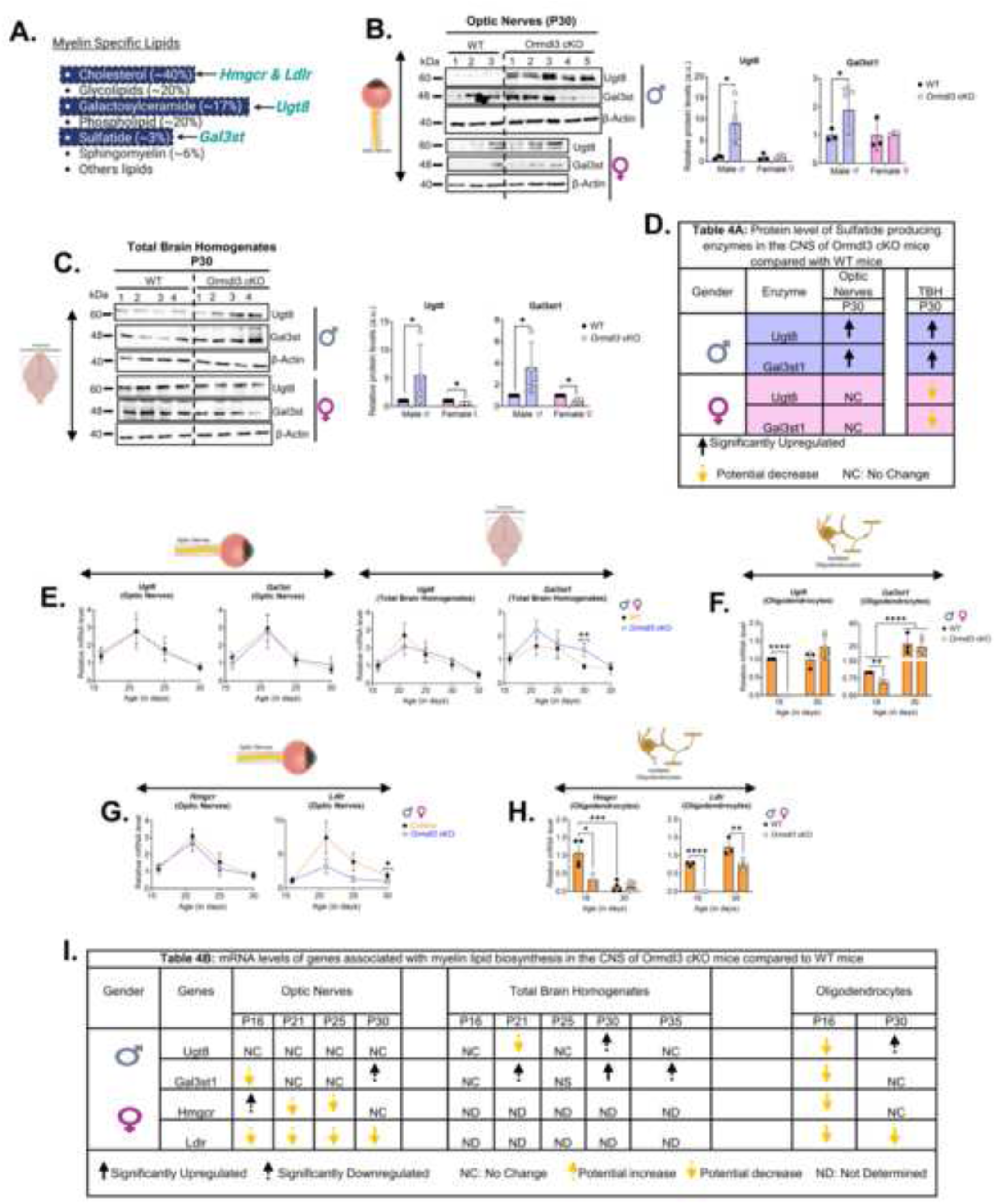
Effect of *Ormdl3* deletion in oligodendrocytes on myelin-specific sphingolipid-synthesizing enzymes and cholesterol synthesis pathway genes. **A.)** Shown is the cartoon representation of different enzymes (teal green colored) that are focused on in this section. Immunoblot analysis of myelin-specific sphingolipid-synthesizing enzymes (UGT8 and GAL3ST1) protein levels. **B.)** UGT8 and GAL3ST1 protein levels at P30 in optic nerve lysates with quantification in WT and *Ormdl3* cKO mice (males and females); **C.)** UGT8 and GAL3ST1 protein levels at P30 in total brain lysates, with quantification, in WT and *Ormdl3* cKO mice (males and females). **D.) Table 4A.** Summary of the results of immunoblotting from panels **A** to **D**. For immunoblotting, 1-2 µg of lysate was loaded into each lane of in-house-made gradient gels. The blots were then probed with UGT8, GAL3ST1, and β-Actin. β-Actin was used as a loading control. Band intensity was then quantified using Quantity One software. Data were normalized to β-Actin and then set relative to WT. Data were shown as the mean ± SD; n=3–4. Shown are the mRNA levels of **E.)** *Ugt8* and *Gal3st1* in optic nerves and the brain tissue from WT and *Ormdl3* cKO mice at P16, P21, P25, and P30 (mixed pool of sexes; **F.)** *Ugt8* and *Gal3st1* in oligodendrocytes isolated from the brains of the WT and *Ormdl3* cKO mice at P16 and P30 (mixed pool of sexes); **G.)** *Hmgcr* and *Ldlr* in optic nerves from WT and *Ormdl3* cKO mice at P16, P21, P25, and P30 (mixed pool of sexes). Shown are the mRNA levels of **H.)** *Hmgcr* and *Ldlr* in oligodendrocytes isolated from the brains of the WT and *Ormdl3* cKO mice at P16 and P30 (mixed pool of sexes). **I.) Table 4B—summary** of the results of real-time PCR from panels **E** to **H**. For mRNA quantification, total mRNA was isolated, and RT-qPCR was performed to quantify mRNA levels as described in the method section. Data was normalized to either *Gapdh* or *Hprt* and set relative to WT (P16). Data were presented as mean ± SD; n = 3–8 (a mixed pool of males and females). Statistical significance was tested by the Student’s two-tailed t-test. *Asterisks* denote significance, p <0.05 = *, p <0.005 = ** and p <0.00005 = ****. *Hprt*: Hypoxanthine phosphoribosyl transferase 1, *Ugt8*: UDP glycosyltransferase 8, *Gal3st*: Galactose-3-O-Sulfotransferase, *Hmgcr*: 3-hydroxy-3-methyl-glutaryl-coenzyme A reductase, and *Ldlr*: low-density lipoprotein receptor.

In the optic nerves at P16, UGT8 protein levels are significantly decreased in cKO mice compared to WT mice (**Sup Fig. 8D**). On the other hand, in the brain at P16, UGT8 protein levels are significantly increased in cKO mice compared to WT mice (**Sup Fig. 8E**). At day 16, mRNA levels of *Ugt8* and *Gal3st1* are not altered in either optic nerves or brain but are significantly downregulated in isolated OLGs in cKO compared with WT (**Fig. 6F**).

Considering the maintenance of cholesterol levels in myelin, suggesting an increase in cholesterol synthesis or uptake to compensate for increased SL levels, we probed the expression of two key elements of cholesterol metabolism, 3-hydroxy-3-methylglutaryl-coenzyme A reductase (Hmgcr), the rate-limiting enzyme in cholesterol synthesis [53], and the low-density lipoprotein receptor (Ldlr) [54], in both optic nerves and brain-derived OLGs at various developmental stages. Surprisingly, the levels of both of these transcripts were either not affected or were decreased in the knockout animals relative to WT at various times of development and in optic nerve and isolated OLGs (**Figs. 6G and H**). This counterintuitive effect is addressed in the discussion.

### 3.7. *Ormdl3* cKO does not alter the number of mature Oligodendrocytes in the Corpus callosum of postnatal day 16 and 30 mice

To assess the effect of *Ormdl3* cKO on OLG differentiation and survival, we analyzed OLG maturation in the corpus callosum of *Ormdl3* cKO mice using immunostaining and confocal microscopy at P16 and P30. The cartoon representation of the process of OLG differentiation is summarized in **Fig. 7A**. The corpus callosum was chosen for this analysis due to its critical role in early brain development, and since the spatiotemporal aspects of OLG differentiation and myelination have been well characterized in this brain region [55–57]. For immunostaining, CC1 is used as a marker to track maturing and myelinating OLGs [58, 59], and Olig2 as a marker to track all OLG lineage cells [60]. Co-localization of Olig2+ with CC1+ positive cells revealed that the number of maturing OLGs is not significantly altered in the corpus callosum of *Ormdl3* cKO compared to WT mice at both ages analyzed (**Figs. 7B-7C**). We examined the number of OLGs in the corpus callosum using coronal sections at P16 and sagittal sections at P30 to provide a holistic overview.

**Figure 7:**
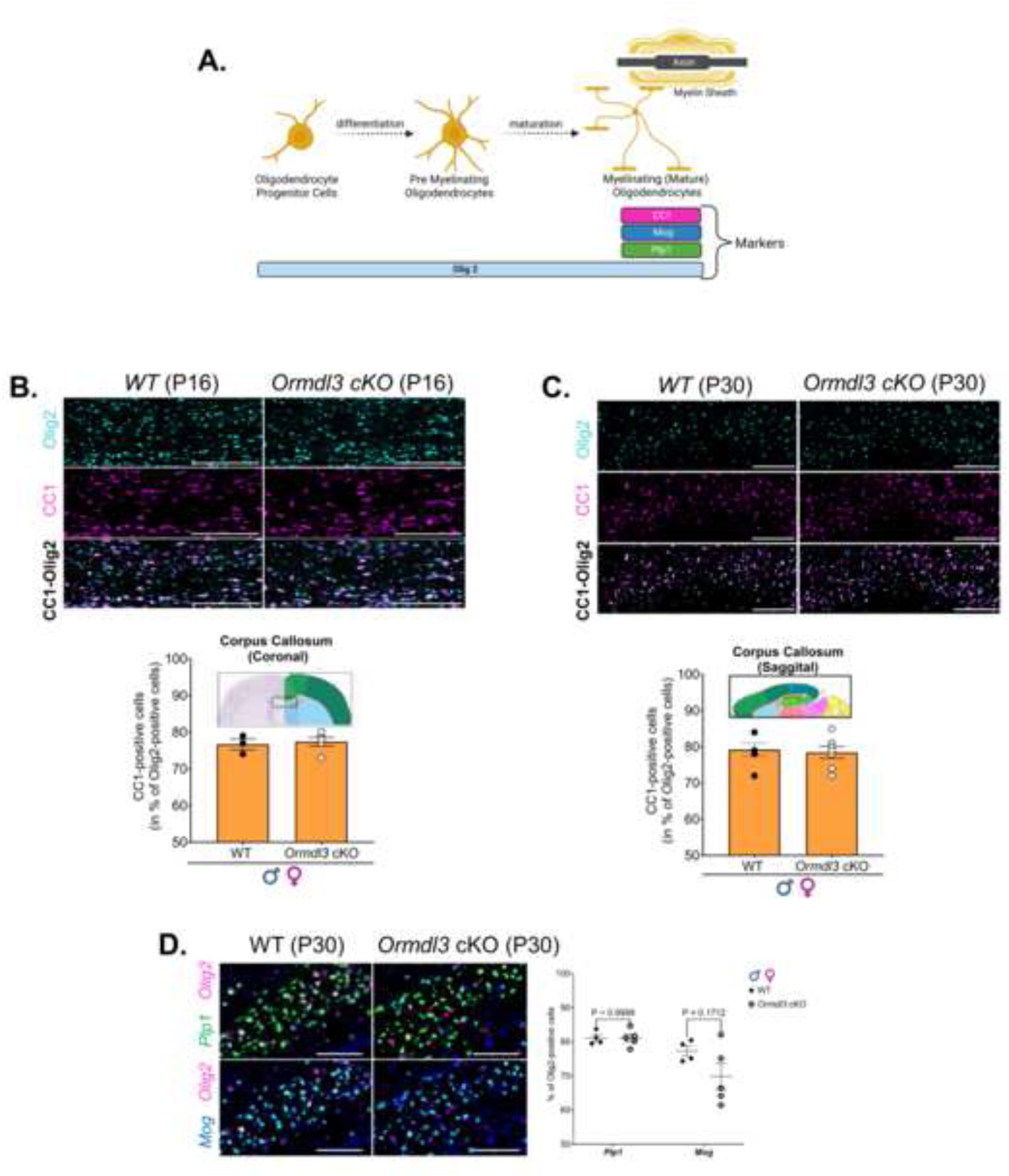
Effect of *Ormdl3* deletion on the number of oligodendrocytes in the corpus callosum. **A.)** Shown is the cartoon representation of a summary of oligodendrocyte differentiation and maturation with associated markers. **B.)** Shown are the immunofluorescence images with quantification of images of the corpus callosum (brain) at P16 (coronal section) and **C.)** P30 (sagittal sections). Immunofluorescence was performed as described in the methods section. In brief, 40 µM sections were prepared and then stained for Olig2 and CC1. For quantification, Olig2- and CC1-positive cells were manually counted using Fiji (ImageJ). Results are graphed as the % of Olig2-positive cells that co-localize with CC2-positive cells. CC1 is a marker for mature and myelinating oligodendrocytes, and Olig2 is a marker for all oligodendrocyte lineage cells. The scale bars for P16 and P30 imaging were set to 200 µm and 500 µm, respectively. Data were presented as mean ± SD; n = 3–6 (a mixed pool of males and females). **D.)** Shown are images after the RNAscope in situ hybridization assay in the sagittal section of the corpus callosum (brain) at P30. RNAscope *in situ* hybridization assay was performed as described in the methods section. In brief, 15-µm sections were prepared and subjected to *in situ* hybridization with probes for Plp1, Mog, and Olig2. For quantification, *Plp1*-, Mog-, and Olig2-positive cells were manually counted using Fiji/ImageJ. Plp1 and Mog are markers for mature and myelinating oligodendrocytes, and Olig2 is a marker for all oligodendrocyte lineage cells. The scale bars were set to 50 µm. Data were presented as mean ± SD; n = 3–6 (a mixed pool of males and females). Statistical significance was tested by the Student’s two-tailed t-test. Olig2: Oligodendrocyte Transcription Factor 2, CC1: anti-adenomatous polyposis coli clone CC1, Plp1: Proteolipid Protein 1, Mog: Myelin Oligodendrocyte Glycoprotein.

To further validate the results obtained by immunohistochemistry, we used RNAscope [59] with probes specific for *Plp1*, *Mog*, and *Olig2* on brain sections from P30 *Ormdl3* cKO and WT mice. PLP1 and MOG serve as established markers for maturing OLGs [61], while Olig2 marks all cells within the OLG lineage [62]. Co-localization of Olig2-positive cells with *Plp1* or *Mog* mRNA revealed that the number of maturing OLGs is not significantly altered in the corpus callosum of *Ormdl3* cKO compared to WT mice (**Fig. 7D**). This observation is consistent with our immunoblot results. Interestingly, *Mog* expression in OLGs has been directly linked to myelination [63, 64], and our RNAscope results point toward a larger variation in *Mog* compared to *Plp1* mRNA levels in *Ormdl3* cKO mice. This observation may indicate compensatory mechanisms specific to myelination rather than to progression through the entire OLG differentiation program. Overall, these results suggest that conditional knockout of *Ormdl3* in OLGs does not significantly affect the timing of OLG differentiation and support the idea that OLGs lacking *Ormdl3* alter their biology to enhance the production of myelin-specific lipids and proteins.

### 3.8. *Ormdl3* cKO in CNS Oligodendrocytes Downregulates mRNA levels of *Sptlc1*, *Sptlc2*, and *Sptssa* in postnatal day 16 mice

To elucidate the effect of *Ormdl3* cKO on other SPT subunits, we measured protein levels of the major SPT subunit SPTLC1 at P16 and mRNA levels of *Ormdl* isoforms (*Ormdl1*, *Ormdl2*, and *Ormdl3*), the major SPT subunits (*Sptlc1*, *Sptlc2*), and the small subunit (*Sptssa*) in brain-derived OLGs from P16 and P30 mice. The cartoon representation of the SPT/ORMDL complex with its subunits is shown in **Fig. 8A**. SPTLC1 protein levels are not affected by *Ormdl3* cKO in OLGs at P16 (**Fig. 8B**). In contrast, there are significant changes in mRNA levels of the SPT subunits at P16. The findings of this section are summarized in **Fig. 8E** (**Table 5**) for clarity. As expected, *Ormdl3* mRNA levels were undetectable at both P16 and P30 (**Fig. 8C**). At P16, *Ormdl1* mRNA levels were significantly downregulated in cKO mice compared to WT mice. Conversely, *Ormdl2* levels were upregulated in cKO mice, likely as a compensatory response to the loss of *Ormdl3*. By P30, the mRNA levels of both *Ormdl1* and *Ormdl2* in cKO mice returned to levels comparable to those of WT mice. The mRNA levels of *Sptlc1*, *Sptlc2*, and *Sptssa* in cKO mice showed a trend similar to that of *Ormdl1* mRNA levels at P16, indicating downregulation compared with WT mice (**Fig. 8D**). By P30, the mRNA levels of all three genes in cKO mice returned to levels comparable to those in WT mice. We observed that the mRNA levels of all SPT subunit genes were significantly downregulated at P30 in both WT and *Ormdl3* cKO mice compared to P16. The implication of this finding is discussed in detail in the discussion section.

**Figure 8:**
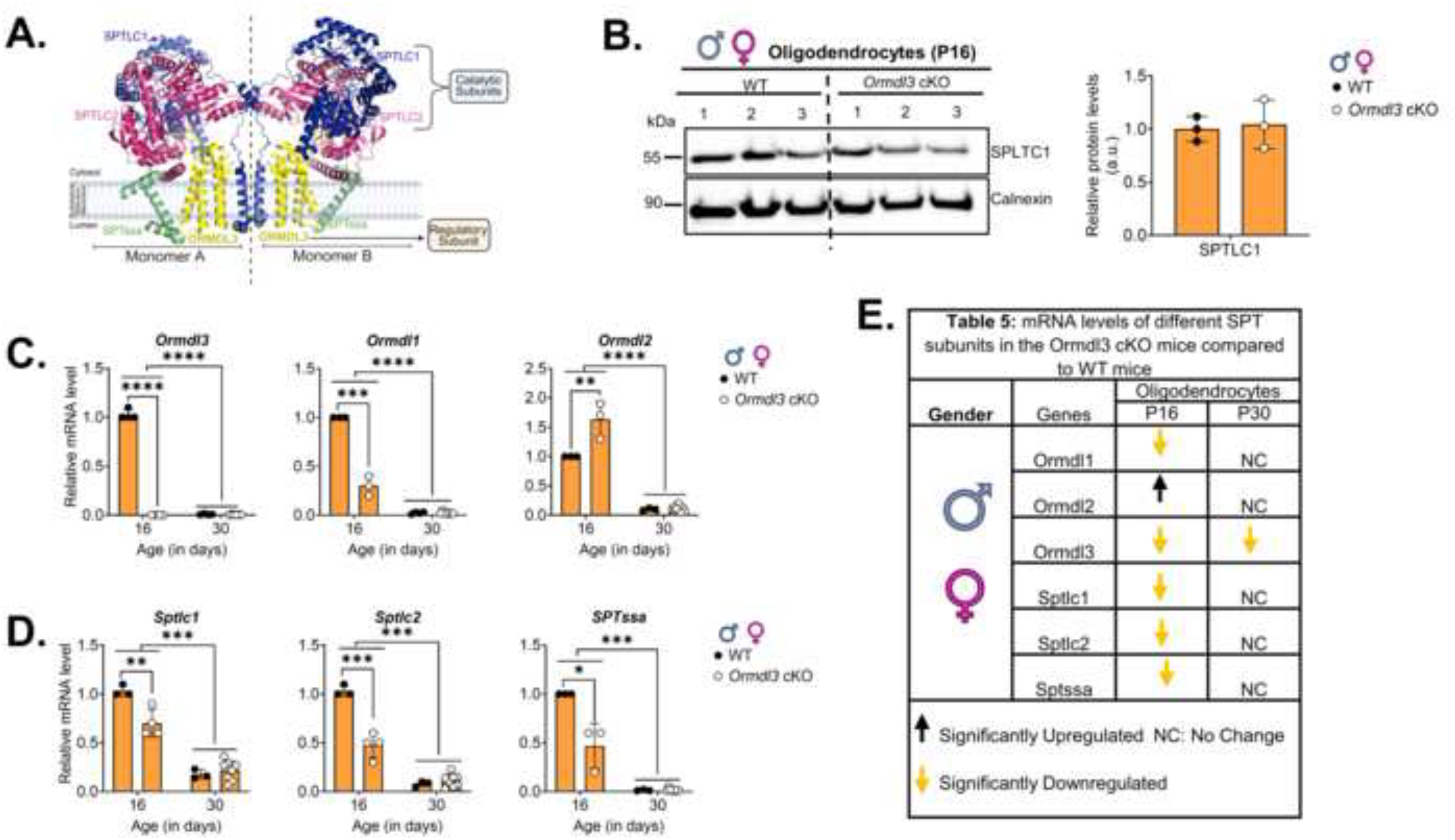
Effect of *Ormdl3* deletion on the other subunits of the serine palmitoyltransferase enzyme. **A.)** Cartoon representation of SPT/ORMDL complex (PDB:7K0n). SPTLC1 is shown in blue, SPTLC2 is shown in pink, SPTssa is shown in cyan, and ORMDL is shown in yellow. Immunoblot analysis of **B.)** SPTLC1 and Calnexin at P16 in isolated oligodendrocytes with quantification. In brief, oligodendrocytes isolated from brains were resuspended in 2X lamellae buffer. Samples were loaded into each lane of in-house-made gradient gels. The blots were then probed with SPTLC1. Calnexin was used as a loading control. The band intensity was then quantified by using Quantity One software. Data were normalized to β-actin and then set relative to WT. Shown are the mRNA levels of **C.)** *Ormdl3*, *Ormdl1*, and *Ormdl2*, **D.)** *Sptlc1*, *Sptlc2*, and *Sptssa* in oligodendrocytes isolated from WT and *Ormdl3* cKO mice at P16 and P30. Total mRNA was isolated, and RT-qPCR was performed to quantify mRNA levels. Data was normalized to *Gapdh* and set relative to WT (P16). Data were presented as mean ± SD; n = 3–8 (a mixed pool of males and females). **E.) Table 5**. Summary of the results of real-time PCR from panels **A** and **B**. Data were shown as the mean ± SD; n = 3–4. *Asterisks* denote significance, p <0.0005 = ***, p <0.00005 = ****.

### 3.9. *Ormdl3* cKO Regulation of Myelin Genes in Mouse Brain is Recapitulated *in vitro* in cultured Rat Oligodendrocytes

To validate our in vivo mouse results, we investigated the effect of *Ormdl3* deletion on SPT subunits and myelin genes in primary rat OLGs. Primary rat OLGs were isolated from P8 rat pups, and *Ormdl3* was transiently depleted using a siRNA knockdown approach. As expected, *Ormdl3* mRNA levels were significantly downregulated in the *Ormdl3* siRNA condition compared with the scramble siRNA condition (control) (**Fig. 9A**). Consistent with our observations in mouse OLGs isolated from WT and cKO animals, the mRNA levels of *Ormdl* and other SPT subunit genes were significantly downregulated in *Ormdl3*-knockdown cells compared with cells treated with scrambled siRNA. The effect of downregulation of *Ormdl1*, *Ormdl2*, and *Sptlc1* mRNA levels was also observed at the protein level. Total ORMDL and SPTLC1 protein levels were significantly downregulated in *Ormdl3*-knockdown cells relative to scrambled siRNA-treated cells (**Fig. 9B**). The mRNA levels of myelin protein genes (*Mbp* and *Mog*) were significantly downregulated in *Ormdl3*-knockdown cells compared to cells treated with scrambled siRNA (**Fig. 9C**). However, consistent with our observations in total brain at day 35, MOG protein levels were elevated by ORMDL3 knockdown despite decreased mRNA levels (**Fig. 9D**). Additionally, *Ugt8* and *Gal3st1* mRNA levels and UGT8 and GAL3ST1 protein levels were significantly decreased in *Ormdl3*-knockdown cells compared to control (**Fig. 9E-9F**). *Hmgcr* and *Ldlr* were also significantly downregulated in *Ormdl3*-knockdown cells compared to control (**Fig. 9G**). These findings align with our observations in the mouse brain. This confirms that the effects we observe in the mouse brain are due to *Ormdl3* deletion and that the results can be replicated in vitro.

**Figure 9:**
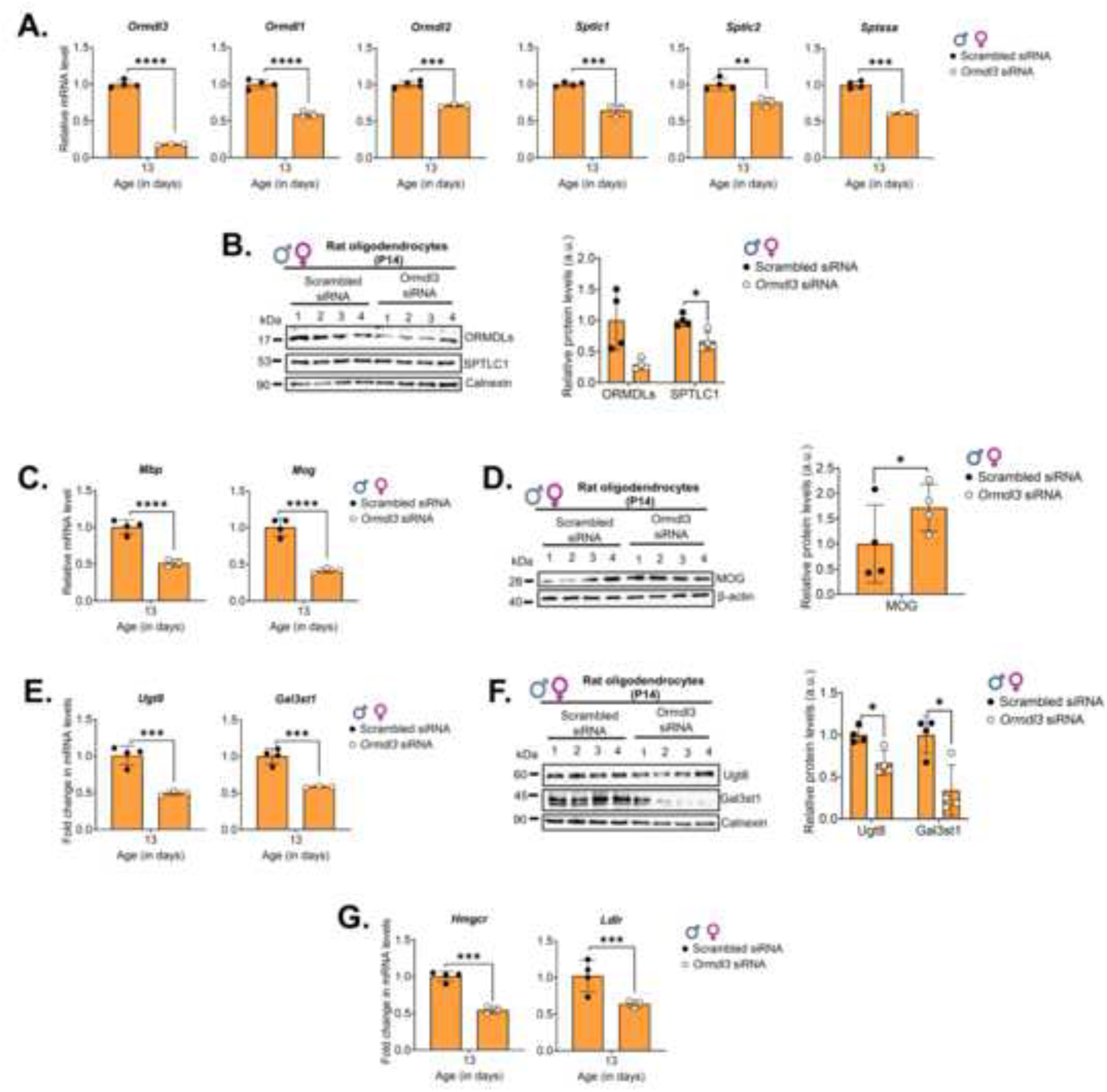
Effect of transient depletion of *Ormdl3* in rat oligodendrocytes on the other subunits of the serine palmitoyltransferase enzyme and myelin genes. Primary oligodendrocytes were isolated from postnatal day 8 (P8) rats and transfected with either scramble siRNA (control) or siRNA targeting *Ormdl3*, as described in the Methods section. **A.)** Shown are the mRNA levels of *Ormdl1*, *Ormdl2*, *Ormdl3*, *Sptlc1*, *Sptlc2*, and *Sptssa* in control and *Ormdl3*-*knockdown* cells. **B.)** Immunoblot analysis of SPTLC1 and ORMDL, with quantification, in control and *Ormdl3*-*knockdown* cells. **C.)** Shown are the mRNA levels of *Mbp* and *Mog,* and **D.)** MOG protein levels with quantification, in control and *Ormdl3*-*knockdown* cells. **E.)** Shown are the mRNA levels of *Ugt8* and *Gal3st,* and **F.)** UGT8 and GAL3ST protein levels with quantification in control and *Ormdl3*-*knockdown* cells. **G.)** Shown are the mRNA levels of *Hmgcr* and *Ldlr* in control and *Ormdl3*-*knockdown* cells. For mRNA measurements, total RNA was isolated from transfected oligodendrocytes (four days after transfection), and RT-qPCR was performed to quantify mRNA levels. Data were normalized to *Gapdh* and set relative to the control. Data are presented as mean ± SD; n = 3–4. For immunoblot analysis, oligodendrocytes were collected after five days post-transfection. Samples were prepared using 2X-Lamelli Buffer and loaded into each lane of in-house-made gradient gels. The blots were probed with SPTLC1, ORMDL, MOG, UGT8, and GAL3ST1 antibodies. Calnexin and β-actin were used as loading controls. Band intensities were quantified using Quantity One software. Data were normalized to either calnexin or β-actin and set relative to the scramble control. Data are shown as mean ± SD; n = 3–4. *Asterisks* denote significance: p < 0.05 = *, p < 0.0005 = ***, and p < 0.00005 = ****.

## 4. Discussion

Our findings emphasize the pivotal role of SL biosynthesis regulation in OLG function and myelination, as these lipids are highly abundant in the CNS. SLs are essential components of the myelin sheath, whose high lipid content requires precise metabolic control during myelin biogenesis [12, 65]. Although the biochemical composition of myelin has been well characterized, a significant gap remains in understanding how OLGs coordinate myelin lipid synthesis with myelin lipid production. In particular, the pathways linking SL biosynthesis to myelin protein expression and assembly remain unclear. Here, we address this gap by defining how OLG-specific *Ormdl3* regulation shapes myelin lipid balance and myelin protein abundance during CNS myelination.

### 4.1. Selective Elevation of Long-Chain Myelin Sphingolipids at P35 in *Ormdl3-*deleted Oligodendrocytes

During rodent CNS development, myelin SL production begins early and peaks around P15-P18 [66, 67], after which their levels are maintained from P30 onwards despite ongoing myelin turnover, which is essential for maintaining a healthy myelin population [68]. In our *Ormdl3* cKO mice, myelin SLs showed a modest but coordinated increase, likely because ORMDL3 loss relieves SPT inhibition and increases SL biosynthetic flux, consistent with partial redundancy among ORMDL isoforms [69].

In our mice, SL changes were selective rather than global, with an increase in myelin SLs: sulfatides and sphingomyelin, while other SL species such as ceramides and galactosylceramides remained unchanged. This pattern is notable because ceramide, a central intermediate in the SL biosynthesis pathway, is the precursor for synthesizing complex SLs [70] and galactosylceramides act as precursors for sulfatide production [71]. From P30 to P35, total myelin SL levels remained stable in WT mice, but their composition shifted from shorter-to-longer chain species; this long-chain SL incorporation was further enhanced in the cKO mice during active CNS myelination [66]. Similar patterns were also observed in the total brain homogenates (**Sup Figs 2A-F, 4D-F, 5D-F, 6D-F, 7D-F, and 8D-F**). This suggests that increased SPT activity raises substrate availability while downstream pathways preferentially channel lipids into myelin SLs. Thus, ORMDL3-dependent SPT inhibition likely acts as a feedback brake during high SL biosynthetic demand, and its loss exaggerates SL production in the *Ormdl3* cKO OLGs. Furthermore, the selective increase in sulfatides after *Ormdl3* deletion suggests that downstream myelin SL pathways are responsive to altered SL flux, but not in a simple linear manner. UGT8 and Gal3st showed sex- and region-dependent protein changes, with increases most evident in male cKO optic nerves and brain, whereas females showed little change or reduction. This pattern suggests that male and female OLGs may adapt differently to disrupted SL homeostasis. However, these protein changes are not consistently matched by mRNA levels. In isolated cKO OLGs, *Ugt8* and *Gal3st1* transcripts were reduced at P16 but returned to WT-like levels by P30, suggesting a transient cell-intrinsic adjustment rather than sustained suppression of sulfatide biosynthetic capacity. Because UGT8 and Gal3st act downstream of SPT to generate galactosylceramides and sulfatides, respectively, this pattern argues against a simple model in which increased sulfatides are driven primarily by upregulation of these enzymes. Instead, loss of ORMDL3-mediated SPT inhibition likely increases upstream SL flux, with UGT8 and Gal3st adjusting secondarily through post-transcriptional or compensatory mechanisms.

Consistent with this broader lipid remodeling, increased SL biosynthesis may also affect cholesterol metabolism. Cholesterol and SLs are closely linked: cholesterol can regulate SL production [72] and elevated sphingomyelin can promote cholesterol biosynthesis[73]. In our study, myelin cholesterol increased in cKO mice, whereas mRNA levels for *Hmgcr,* the rate-limiting enzyme in the cholesterol biosynthetic pathway [53, 74], and *Ldlr,* a major cholesterol uptake receptor [54] were reduced. As OLGs also receive astrocyte-derived cholesterol during myelin formation [75, 76], the observed persistent reduction in cholesterol metabolism genes likely reflects a complex response to altered lipid balance rather than a simple decrease in cholesterol demand. Thus, *Ormdl3* loss may secondarily reshape cholesterol homeostasis while altering myelin lipid balance.

### 4.2. *Ormdl3* cKO and its effect on the SPT complex

The increase in myelin SL in *Ormdl3* cKO mice does not appear to result from higher SPT protein abundance. Although SPT subunit transcripts normally decline from P16 to P30 during OLG maturation [6] SPTLC1 protein levels were unchanged in cKO OLGs, suggesting that SPT complex abundance is maintained despite reduced mRNA levels. The compensatory increase in *Ormdl2* expression may partially replace *Ormdl3*, but because ORMDL2 is less sensitive to ceramide-dependent feedback regulation, this shift may permit greater SPT activity during later myelination [18, 19]. The SPT enzyme complex is composed of subunits in a strict 1:1:1:1 stoichiometric ratio (SPTLC1:SPTLC2:SPTSSA:ORMDL), which is essential for enzymatic activity. Importantly, Sptlc1 protein expression determines the stability and abundance of the other subunits [77]. Together, these findings suggest that *Ormdl3* deletion increases SL flux primarily by relaxing SPT inhibition rather than by increasing SPT protein levels, consistent with prior evidence that SPT activity can change independently of SPT protein abundance [6].

### 4.3. Enrichment of Large-Caliber Axons Without g-Ratio Change in Optic Nerves of *Ormdl3* cKO Mice

We assessed the myelin phenotype in optic nerves, which provide a useful system for examining how OLG lipid metabolism influences myelination, owing to their anatomical and functional simplicity relative to the brain [78]. In our myelin ultrastructure study, thicker myelin sheaths, enrichment of larger-caliber axons, and a preserved g-ratio in Ormdl3 cKO mice suggest that increased myelin production is not merely excess membrane deposition but is proportionally matched to axon size. This raises the possibility that enhanced myelin biosynthetic capacity is coordinated with local axonal cues that preserve appropriate myelin thickness relative to axon caliber. Such coordination may involve bidirectional signaling, in which axons influence myelin sheath formation while OLGs provide feedback that supports axon caliber during development [31]. Thus, the structural phenotype is best interpreted as adaptive remodeling of the axon-myelin unit rather than isolated myelin overgrowth. Whether increased myelin synthesis drives axon enlargement or whether larger axons preferentially recruit additional myelin remains unresolved. Further defining the directionality and molecular basis of this interaction will be important for understanding how SL regulates CNS myelin architecture.

### 4.4. Transcriptional and Translational Upregulation of Myelin Protein Genes in *Ormdl3* cKO Mice

Along with the myelin lipid and ultrastructural changes, the increase in myelin proteins in *Ormdl3* cKO mice suggests that altered SL biosynthesis influences broader myelin assembly rather than lipid composition alone. It is well established that myelin assembly and compaction depend on tightly regulated myelin protein levels [4]. While our data hint at an interesting sex-dependent difference in protein abundance in *Ormdl3* cKO mice, further studies may help in deciphering sex differences in susceptibility to demyelinating disease [79, 80]. Our link between SL upregulation and increased myelin protein levels is consistent with prior evidence that reduced sulfatide levels decrease myelin protein abundance in aged mice [81] and with studies showing that depletion of ceramides and C22/C24 chain-length SLs through whole-body or OLG-specific ceramide synthase 2 knockout mice leads to hypomyelination [82, 83]. Although WT mice normally show declining *Mbp, Mog,* and *Plp1* transcripts with aging, consistent with previous reports [6, 84, 85], *Ormdl3* cKO mice showed increased myelin protein levels with region-specific changes in mRNA levels, suggesting both transcriptional and translational regulation. Notably, *Plp1* mRNA levels were reduced in isolated cKO OLGs but not in total brain or optic nerves, likely because whole-tissue measurements average signals from existing myelin and a regionally distinct OLG population. Thus, this OLG-specific reduction may reflect a cell-intrinsic response to altered SL homeostasis that is masked at the tissue level. Together with transient *Ormdl3* knockdown data (**Section 4.6**), these findings suggest that SL availability helps coordinate myelin lipid and protein production through both transcriptional and post-transcriptional mechanisms that remain to be defined.

### 4.5. *Ormdl3* Loss Alters Myelin Biosynthesis Without Changing Oligodendrocyte Numbers and Differentiation

While we observed thicker myelin sheaths in the cKO mice, associated with higher SL production resulting in increased myelin protein levels, OLGs in the corpus callosum of our cKO mice showed no changes in OLG number or maturation at P16 and P30. This interesting phenotype likely reflects altered metabolic activity in differentiated OLGs rather than impaired development. OLGs undergo a complex, tightly regulated differentiation program in which proliferative OLG progenitor cells (OPCs) progress through premyelinating stages before becoming mature myelinating cells [25, 26]. This transition requires coordinated transcriptional, post-transcriptional, metabolic, and membrane-remodeling events that support axon engagement, myelin membrane expansion, and sheath compaction. Thus, our study suggests that ORMDL3-dependent restraint of SPT may tune myelin membrane assembly by regulating SL flux within mature OLGs. The early reduction in myelin- and SL-related transcripts may reflect feedback suppression after increased SPT activity, whereas their later restoration suggests compensation during ongoing myelination. Thus, *Ormdl3* loss shifts the lipid metabolic environment that supports membrane growth, protein incorporation, and axon–myelin proportionality.

### 4.6. Transient *Ormdl3* Downregulation in Rat Oligodendrocytes Recapitulates *In vivo*

#### Transcriptional Suppression of Myelin Biogenesis in *Ormdl3* cKO mice

The above-discussed observations link the *in vivo* phenotype to altered SPT regulation, but they do not distinguish whether the transcriptional changes arise directly from intrinsic OLG *Ormdl3* loss. To address this, we transiently depleted *Ormdl3* in isolated rat OLGs. Similar to the mouse cKO, *Ormdl3* knockdown broadly reduced the expression of transcripts encoding ORMDL/SPT subunits, myelin proteins, and enzymes involved in sulfatide and cholesterol metabolism. The downregulation of *Ormdl3* is also accompanied by reduced protein levels, suggesting that ORMDL3 is a prominent ORMDL isoform present. The downregulation of SPTLC1 protein levels suggests a transient response to Sptlc1 mRNA depletion. A similar observation was made for the UGT8 and Gal3ST proteins: downregulation of mRNA levels was reflected in protein levels, further validating that the *Ormdl3* knockdown transiently alters genes and enzymes associated with myelin synthesis. However, reduced transcript abundance was not uniformly matched at the protein level for myelin protein; Mog protein increased despite lower *Mog* mRNA, supporting the idea that post-transcriptional mechanisms contribute to myelin protein regulation when SL homeostasis is disrupted. Thus, the rat OLG model recapitulates key effects of *Ormdl3* loss and provides a tractable system for dissecting how altered SL flux coordinates myelin lipid metabolism with protein production.

## 5. Conclusion

Together, our findings identify OLG ORMDL3 as a key regulator of CNS myelin composition and architecture. Loss of *Ormdl3* selectively increased long-chain myelin SL, elevated major myelin proteins, and altered optic nerve ultrastructure without changing OLG number or differentiation. These changes are best explained by reduced ORMDL3-dependent inhibition of SPT, thereby increasing SL flux and shifting the metabolic environment in which mature OLGs assemble myelin. The preserved g-ratio despite thicker myelin and larger-caliber axons further suggests that lipid-driven myelin remodeling remains coordinated with axon size rather than causing unbalanced membrane accumulation. Thus, ORMDL3 acts as a metabolic brake that helps synchronize SL production with myelin protein abundance and axon–myelin proportionality during development. By defining this OLG-intrinsic regulatory mechanism, this study provides a foundation for understanding how disrupted SL homeostasis may contribute to demyelinating and dysmyelinating diseases.

## 6. Acknowledgments

We are thankful to Dr. Klaus-Armin Nave from the Max Planck Institute in Germany for generously agreeing to use the Cnp-Cre mouse line. The VCU Massey Cancer Center Lipidomics and Metabolomics Shared Resource provided services in support of the research project, supported in part by NIH-NCI Cancer Center Support Grant P30 CA016059. Special thanks to Dr. Jeremy Allegood for his advice and support in lipidomic analysis. For the generation of RNAscope and confocal imaging data, support was provided through the VCU Massey Comprehensive Cancer Center Microscopy and TDAAC (Tissue and Data Acquisition and Analysis) Shared Resources, supported in part by funding from the NIH-NCI Cancer Center Support Grant P30 CA016059. All the cartoon images are “Created with BioRender.com”. This work was supported by R21NS120128 awarded to BW.

## 7. Disclosures regarding the Use of AI

This manuscript is written without the assistance of any AI tools. Grammarly software has been used to correct grammatical errors.

## 8. Conflict of Interest

The authors declare no conflict of interest.

## 9. Authors’ Contributions

**UM**: Writing – review & editing, Writing – original draft, visualization, validation, supervision, project administration, methodology, investigation, formal analysis, data curation, conceptualization, generated data for all the figures except data for figure 7, **KM**: guidance for mouse handling and maintenance, **RF**: quantification of EM images, **FF**: cholesterol assay, **SS**: helped with myelin isolation, collecting brains, editing manuscript, **CSB:** guidance and helped with oligodendrocyte and myelin isolation, **TK**: guidance for mice handling, **BF**: supervised and generated, analyzed, and graphed all data for RNAscope and immunostaining, **FSA**: immunostaining, **SM:** Sectioning of brains for RNAscope and immunostaining, **JD**: guidance and helped with EM imaging, quantification, and analysis, **BW:** Writing – review & editing, Validation, Supervision, Resources, Project administration, Methodology, Investigation, Funding acquisition, Formal analysis, Data curation, Conceptualization, and Cholesterol assay.

## Abbreviations

cKO: Conditional knockout
CNPase: 2’,3’-cyclic nucleotide 3-phosphodiesterase
CNS: Central nervous system
CoA: Coenzyme A
CNP: 2’,3’-cyclic nucleotide 3-phosphodiesterase
ER: Endoplasmic reticulum
Gal3st1: Galactose-3-O-sulfotransferase 1
GalCer: Galactosylceramide
GFAP: Glial fibrillary acidic protein
GlcCer: Glucosylceramide
Hmgcr: 3-hydroxy-3-methylglutaryl-coenzyme A reductase
KO: Knockout
Ldlr: Low-density lipoprotein receptor
MBP: Myelin basic protein
MOG: Myelin oligodendrocyte glycoprotein
mRNA: Messenger RNA
PCR: Polymerase chain reaction
PLP: Proteolipid protein
PNS: Peripheral nervous system
RNA: Ribonucleic acid
Sa: Sphinganine
Sa1P: Sphinganine 1-phosphate
siRNA: Small interfering RNA
SL: Sphingolipid
SM: Sphingomyelin
So: Sphingosine
So1P: Sphingosine 1-phosphate
SPTLC1/2/3: Serine palmitoyltransferase long-chain base subunits 1, 2, and 3
SPTssa/SPTssb: Small subunits of serine palmitoyltransferase a and b
TBH: Total brain homogenate Ugt8
UDP: glycosyltransferase 8
WT: Wild type

## Conflict of Interest

The authors declare that they have no known competing financial interests or personal relationships that could have appeared to influence the work reported in this paper.

## Notes

### Competing Interest Statement

The authors have declared no competing interest.

