## Supplementary Figure Legends for "ORMDL3-mediated SPT regulation Coordinates Myelin Sphingolipid and Protein Synthesis in Oligodendrocytes"

**Supplementary Figure 1: Levels of different acyl chain length Sulfatides (non-Hydroxylated and hydroxylated) in myelin from postnatal day 30 and 35 mice**

**A.)** PCR amplification of wild-type *Ormdl3* in total genomic DNA was performed using brain lysates extracted from postnatal day 35 WT and *Ormdl3* cKO mice. The steady-state levels of **B.)** different acyl chain lengths of non-hydroxylated sulfatide species in the myelin isolated from P30 mouse brains from both male and female mice, respectively; **C.)** different acyl chain lengths of non-hydroxylated sulfatide species in the myelin from P35 WT and *Ormdl3* cKO mice, male and female, respectively. **D.)** Different acyl chain lengths of hydroxylated sulfatide species in the myelin from P30 in WT and *Ormdl3* cKO mice, male and female, respectively; **E.)** Different acyl chain lengths of hydroxylated sulfatide species in the myelin from P35 in WT and *Ormdl3* cKO mice, male and female, respectively. Sulfatide levels were measured by Mass Spectrometry as described in the methods section. The lipidomic data are graphed as picomoles of lipids per 100 µg of protein. Shown are the mean ± SD, n=3-4. Statistical significance was tested by the Student’s two-tailed *t*-test. *Asterisks* denote significance, *p* <0.05 = *. Different acyl chain lengths measured are C14:0, C16:0, C18:1, C18:0, C20:0, C22:0, C24:1, C24:0, C26:1, and C26:0

**Supplementary Figure 2: Levels of different acyl chain length Sulfatides (non-Hydroxylated and hydroxylated) in total brain homogenates from postnatal day 30 and 35 mice**

The steady-state levels of **A.)** total sulfatides (non-hydroxylated) in the total brain homogenates of P30 and P35 WT and *Ormdl3* cKO mice; **B.)** different acyl chain lengths of non-hydroxylated sulfatide species in P30 mouse brain from both male and female mice, respectively; **C.)** different acyl chain lengths of non-hydroxylated sulfatide species in P35 mouse brain from both male and female mice, respectively. The steady-state levels of **D.)** total sulfatides (hydroxylated) in the total brain homogenates of P30 and P35 WT and *Ormdl3* cKO mice; **E.)** different acyl chain lengths of hydroxylated sulfatide species in the total brain homogenates of P30 mice, both male and female mice, respectively; **F.)** different acyl chain lengths of hydroxylated sulfatide species in the total brain homogenates of P35 mice, both male and female mice, respectively. Sulfatide levels were measured by Mass Spectrometry. The lipidomic data are graphed as picomoles of lipids per 100 µg of protein. Shown are the mean ± SD, n=3-4. Statistical significance was tested by the Student’s two-tailed *t*-test. *Asterisks* denote significance, *p* <0.05 = *. Different acyl chain lengths measured are C14:0, C16:0, C18:1, C18:0, C20:0, C22:0, C24:1, C24:0, C26:1, and C26:0

**Supplementary Figure 3: Levels of sphingoid long-chain bases in both tissue lysates of the total cerebral hemispheres and myelin from postnatal day 30 and 35 mice**

The steady-state levels of **A.)** dihydrosphingosine in the myelin of P30 and P35 WT and *Ormdl3* cKO mice; **B.) – D.)** sphingosine, sphingosine-1-phosphate, and dihydrosphingosine in the total brain homogenates from WT and *Ormdl3* cKO mice, at P30 and P35, male and female, respectively. The sphingolipid levels were measured by Mass Spectrometry. The lipidomic data are graphed as picomoles of lipids per 100 µg of protein. Shown are the mean ± SD, n=3-4. Statistical significance was tested by the Student’s two-tailed *t*-test. *Asterisks* denote significance, *p* <0.05 = *, *p* <0.005 = ** and *p* <0.0005 = ***.

**Supplementary Figure 4: Levels of sphingomyelin in both total brain lysates and myelin from postnatal day 30 and 35 mice**

The steady-state levels of **A.)** total sphingomyelin in myelin of P30 and P35 WT and *Ormdl3* cKO mice; **B.)** different acyl chain lengths of sphingomyelin species in P30 mice brain from both male and female mice, respectively; **C.)** different acyl chain lengths of sphingomyelin species in P35 mice brain from both male and female mice, respectively. The steady-state levels of **D.)** total sphingomyelin in total brain homogenates of P30 and P35 WT and *Ormdl3* cKO mice; **E.)** different acyl chain lengths of sphingomyelin species in total brain homogenates of P30 mice, both male and female mice, respectively; **F.)** different acyl chain lengths of sphingomyelin species in total brain homogenates of P35 mice, both male and female mice, respectively. Sphingomyelin levels were measured by Mass Spectrometry. The lipidomic data are graphed as picomoles of lipids per 100 µg of protein. Shown are the mean ± SD, n=3-4. Statistical significance was tested by the Student’s two-tailed *t*-test. *Asterisks* denote significance, *p* <0.05 = * and *p* <0.005 = **. Different acyl chain lengths measured are C14:0, C16:0, C18:1, C18:0, C20:0, C22:0, C24:1, C24:0, C26:1, and C26:0 .

**Supplementary Figure 5: Levels of ceramide in both total brain lysates and myelin from postnatal day 30 and 35 mice**

The steady-state levels of **A.)** total ceramide in myelin of P30 and P35 WT and *Ormdl3* cKO mice; **B.)** different acyl chain lengths of ceramide species in P30 mice brain from both male and female mice, respectively; **C.)** different acyl chain lengths of ceramide species in P35 mice brain from both male and female mice, respectively. The steady-state levels of **D.)** total ceramide in total brain homogenates of P30 and P35 WT and *Ormdl3* cKO mice; **E.)** different acyl chain lengths of ceramide species in total brain homogenates of P30 mice, both male and female mice, respectively; **F.)** different acyl chain lengths of ceramide species in total brain homogenates of P35 mice, both male and female mice, respectively. Ceramide levels were measured by Mass Spectrometry. The lipidomic data are graphed as picomoles of lipids per 100 µg of protein. Shown are the mean ± SD, n=3-4. Statistical significance was tested by the Student’s two-tailed *t*-test. *Asterisks* denote significance, *p* <0.05 = * and *p* <0.005 = **. Different acyl chain lengths measured are C14:0, C16:0, C18:1, C18:0, C20:0, C22:0, C24:1, C24:0, C26:1, and C26:0

**Supplementary Figure 6: Levels of glucosylceramides in both total brain lysates and myelin from postnatal day 30 and 35 mice**

The steady-state levels of **A.)** total glucosylceramides in myelin of P30 and P35 WT and *Ormdl3* cKO mice; **B.)** different acyl chain lengths of glucosylceramide species in P30 mice brain from both male and female mice, respectively; **C.)** different acyl chain lengths of glucosylceramide species in P35 mice brain from both male and female mice, respectively. The steady-state levels of **D.)** total glucosylceramides in total brain homogenates of P30 and P35 WT and *Ormdl3* cKO mice; **E.)** different acyl chain lengths of glucosylceramide species in total brain homogenates of P30 mice, both male and female mice, respectively; **F.)** different acyl chain lengths of glucosylceramide species in total brain homogenates of P35 mice, both male and female mice, respectively. Glucosylceramide levels were measured by Mass Spectrometry. The lipidomic data are graphed as picomoles of lipids per 100 µg of protein. Shown are the mean ± SD, n=3-4. Statistical significance was tested by the Student’s two-tailed *t*-test. *Asterisks* denote significance, *p* <0.05 = * and *p* <0.005 = **. Different acyl chain lengths measured are C14:0, C16:0, C18:1, C18:0, C20:0, C22:0, C24:1, C24:0, C26:1, and C26:0

**Supplementary Figure 7: Levels of galactosylceramides in both total brain lysates and myelin from postnatal day 30 and 35 mice**

The steady-state levels of **A.)** total galactosylceramides in myelin of P30 and P35 WT and *Ormdl3* cKO mice; **B.)** different acyl chain lengths of galactosylceramide species in P30 mice brain from both male and female mice, respectively; **C.)** different acyl chain lengths of galactosylceramide species in P35 mice brain from both male and female mice, respectively. The steady-state levels of **D.)** total galactosylceramides in total brain homogenates of P30 and P35 WT and *Ormdl3* cKO mice; **E.)** different acyl chain lengths of galactosylceramide species in total brain homogenates of P30 mice, both male and female mice, respectively; **F.)** different acyl chain lengths of galactosylceramide species in total brain homogenates of P35 mice, both male and female mice, respectively. Galactosylceramide levels were measured by Mass Spectrometry. The lipidomic data are graphed as picomoles of lipids per 100 µg of protein. Shown are the mean ± SD, n=3-4. Statistical significance was tested by the Student’s two-tailed *t*-test. *Asterisks* denote significance, *p* <0.05 = * and *p* <0.005 = **. Different acyl chain lengths measured are C14:0, C16:0, C18:1, C18:0, C20:0, C22:0, C24:1, C24:0, C26:1, and C26:0

**Supplementary Figure 8: Levels of Mylein proteins, Ugt8, and Gal3st1 enzyme levels in optic nerves and brain.**

Immunoblot analysis of myelin proteins: **A.)** at P30 in total brain lysates with quantification of MBP, MOG, and PLP in both genders, WT and *Ormdl3* cKO mice; **B.)** at P16 in optic nerve lysates with quantification of MBP, MOG, and PLP; **C.)** at P16 in total brain lysates with quantification of MBP, MOG, and PLP in mixed pools of both genders, WT and *Ormdl3* cKO mice, respectively. **D.)** Immunoblot analysis of Ugt8 in P16 optic nerves with quantification and **E.)** UGT8 in total brain homogenates at P16, with quantification in mixed pools of both genders (WT and *Ormdl3* cKO).

In brief, 1-2 µg of lysate was loaded into each lane of in-house-made gradient gels as described in the methods section. The blots were then probed with MBP, MOG, PLP, GAPDH, and β-actin.  GAPDH and β-actin were used as loading controls. The band intensity was then quantified by using Quantity One software. Data were normalized to GAPDH or β-actin and then set relative to WT. Data were shown as the mean ± SD; n=4. Statistical significance was tested by the Student’s two-tailed t-test. *Asterisks* denote significance, p <0.05 =*. Here, we refer to the brain as the cerebral hemisphere, excluding the cerebellum.
