## Supplementary Tables for "ORMDL3-mediated SPT regulation Coordinates Myelin Sphingolipid and Protein Synthesis in Oligodendrocytes"

| **Supplementary Table 1: Primers used for Genotyping** | | |
| --- | --- | --- |
| **Gene** | **Forward Primer** | **Reverse Primer** |
| *Cnp*_WT | 5’GCCTTCAAACTGTCCATCTCCTC3’ | 5’CCCAGCCCTTTTATTACCAC3’ |
| *Cnp*_Cre | 5’CATAGCCTGAAGAACGAGA3’ | 5’CCCAGCCCTTTTATTACCAC3’ |
| *mOrmdl3* | 5’AGTTCCAGGGCAGGCTTT3’ | 5’ACTGCCGCTCTGCAAAAGA3’ |

| **Supplementary Table 2: Primers used for verification of Ormdl3 cKO at genomic DNA level** | | |
| --- | --- | --- |
| **Gene** | **Forward Primer** | **Reverse Primer** |
| *mOrmdl3*-Set 1 | 5’ATGACAGCCTGGTGTTAGT3’ | 5’GCTCAGAACTCCCCTCTC3’ |
| *mOrmdl3*-Set 1 | 5’AGTGGACAGTGGTGTGGAG3’ | 5’CAGCATCGTTCTGTGGGTA3’ |

| **Supplementary Table 3: qPCR probes for mouse genes (SYBR green)** | | |
| --- | --- | --- |
| **Gene** | **Forward Primer** | **Reverse Primer** |
| *Mbp* | 5’ GTGACACCTCGACACCCCCTCCA3’ | 5’ GCTAAATCTGCTGAGGGACAGGCC3’ |
| *Mog* | 5’ ATCTGCTACAACTGGCTGCAC3’ | 5’GGGAAATCCCCAAGGACCTGC 3’ |
| *Plp1* | 5’ CCACACTAGTTTCCCTGCTCACCT3’ | 5’ GGTGCCTCGGCCCATGAGTT3’ |
| *Hmgcr* | 5’ CACATTCACTCTTGACGCTCT3’ | 5’ ACTGACATGCAGCCGAAG3’ |
| *Ldlr* | 5’ AGACCCAGAGCCATCGTAG3’ | 5’ GATGTCCACACATTCAAACC3’ |
| *Gapdh* | 5’AATGGTGAAGGTCGGTGTG 3’ | 5’GTGGAGTCATACTGGAACATGTAG 3’ |
| *Hprt* | 5’ CCCCAAAATGGTTAAGGTTGC3’ | 5’AACAAAGTCTGGCCTGTATCC 3’ |

| **Supplementary Table 4: qPCR probes for mouse genes (TaqMan probes)** | | | |
| --- | --- | --- | --- |
| **Gene** | **Forward Primer** | **Reverse Primer** | **Probe** |
| *Ormdl1* | 5’CTCTTGCCTGGACCTTAACC3’ | 5’CAGCTGTTCCCAGTGAGTTAG3’ | 5’TTGCTCTAC/ZEN/CCTGATCCGGAGTCT3’ |
| *Ormdl2* | 5’CACTTCCTGGAGACCACAG3’ | 5’AGTCCAACTCAGTCTCCTCAT3’ | 5’CCTGTTCCC/ZEN/CAGCTGTCTCCTGA3’ |
| *Ormdl3* | 5’CTGTCGTCTGGACCCTCA3’ | 5’TGAACTGGACCCCGTAGT3’ | 5’TGTACATGC/ZEN/CCAAGGTTGTGGATAAGG3’ |
| *Sptlc1* | 5’GCCATTCCTGCGTACTCTAAG3’ | 5’CTTGATGCCTGTAATCCTTTCTG3’ | 5’CATCATCTT/ZEN/TGTGGACAGTGCGGC3’ |
| *Sptlc2* | 5’AGGCGTGGTAGATTACTTTGG3’ | 5’ACAGCACTGTGAGAATGTGT3’ | 5’TCGGAGGCA/ZEN/AGAAGGAGCTGATAGA3’ |
| *Sptssa* | 5’ACTGCGCTCTACATGCTG3’ | 5’CAAAGTAATGCAGAATAGCCATGA3’ | 5’CGTGTTCAA/ZEN/TTCGATGCTGGTTTCCG3’ |
| *Ugt8* | 5’GCAGAGGGCTCAGAAGTTATC3’ | 5’ACTACATTCTTCGCCACGAC3’ | 5’TTCATAAGG/ZEN/ATCAACCCGGCCACC3’ |
| *Gal3st1* | 5’CATCAGTGGTTTCCTGAGATGA3’ | 5’CCTATTGCTGCTGTACTCCTATG3’ | 5’TACTGCCGA/ZEN/AGAAGCCCTGCAA3’ |
| *Hrpt* | 5’CCCCAAAATGGTTAAGGTTGC3’ | 5’AACAAAGTCTGGCCTGTATCC3’ | 5’CCTGCTGGT/ZEN/GAAAAGGACCTCTCGAA3’ |
| *Gapdh* | 5’AATGGTGAAGGTCGGTGTG3’ | 5’GTGGAGTCATACTGGAACATGTAG3’ | 5’TGCAATGG/ZEN/CAGCCCTGGTG3’ |

| **Supplementary Table 5: qPCR probes for rat genes (TaqMan probes)** | | | |
| --- | --- | --- | --- |
| **Gene** | **Forward Primer** | **Reverse Primer** | **Probe** |
| *Ormdl1* | 5’ GGCCTGCTTCACATTGTATTT3’ | 5’ CGACCTCACCAGCAAGAATA3’ | 5’ AGTGTTCCT/ZEN/GTTGCCTGGACCTT3’ |
| *Ormdl2* | 5’ CTCCTCAGCATCCCTTTCTTC3’ | 5’ GCGTCCCTTTGACTGTATGT3’ | 5’ AGCATTCCT/ZEN/GTTGTCTGGACCCTG3’ |
| *Ormdl3* | 5’ ACTGGGAGCAGATGGACTAT3’ | 5’ GGACTTGGTCATACTTGGTGTAG3’ | 5’ TGTGCTGTA/ZEN/CTTCCTCACCAGCTT3’ |
| *Sptlc1* | 5’ GAACCATCCTGCTCTCAACTAC3’ | 5’ CCAGCAACCCGAGGAAATTA3’ | 5’ ACCCATAAC/ZEN/ATCGTGGTGAACGGA3’ |
| *Sptlc2* | 5’ TGGTGCCTTTGATGCTCTAC3’ | 5’ GGAAATCCCACCACAACTACA3’ | 5’ CGCCTTTGG/ZEN/GAGAGAGATGCTGAA3’ |
| *Sptssa* | 5’ GAAGCAGATGTCCTGGTTCTAC3’ | 5’ CACAGAAACCAGCATCGAATTG3’ | 5’ TTGCTGGTC/ZEN/ACTGCGCTCTACAT3’ |
| *Ugt8* | 5’GCAGAGGGCTCAGAAGTTATC3’ | 5’ACTACATTCTTCGCCACGAC3’ | 5’TTCATAAGG/ZEN/ATCAACCCGGCCACC3’ |
| *Gal3st1* | 5’CATCAGTGGTTTCCTGAGATGA3’ | 5’CCTATTGCTGCTGTACTCCTATG3’ | 5’TACTGCCGA/ZEN/AGAAGCCCTGCAA3’ |
| *Hrpt* | 5’ GCTTTTCCACTTTCGCTGATG3’ | 5’ GGTGAAAAGGACCTCTCGAAG3’ | 5’TGGATACA/ZEN/GCCAGACTTTGTTGGATT 3’ |
| *Mbp* | 5’ CCTCTCCCTCAGCAGATTTAG3’ | 5’ TCCCTTGTGAGCCGATTTATAG3’ | 5’ CAGAAGCCA/ZEN/GGATTTGGCTACGGA3’ |
| *Mog* | 5’ GCCGTGGAGTTGAAAGTAGAA3’ | 5’ GCTGCAGGAAGAGGAATACAA3’ | 5’ TCTCATTGC/ZEN/CCTTGTGCCTATGCT3’ |
| *Hmgcr* | 5’ GCAACCTCTACATCCGTCTC3’ | 5’ TGCAGCTCAGGAAAGAACTC3’ | 5’ CTCGTCACC/ZEN/CTTGGCCCTCATGTTCATCC3’ |
| *Ldlr* | 5’ AGACCCAGAGCCATCGTAG3’ | 5’ CTACACCATTCAAACCCCCTT3’ | 5’ ATCTGTCCA/ZEN/GTACATGAAGCCATGCA3’ |
