## Supplementary Figures for "ORMDL3-mediated SPT regulation Coordinates Myelin Sphingolipid and Protein Synthesis in Oligodendrocytes"

### Slide 1
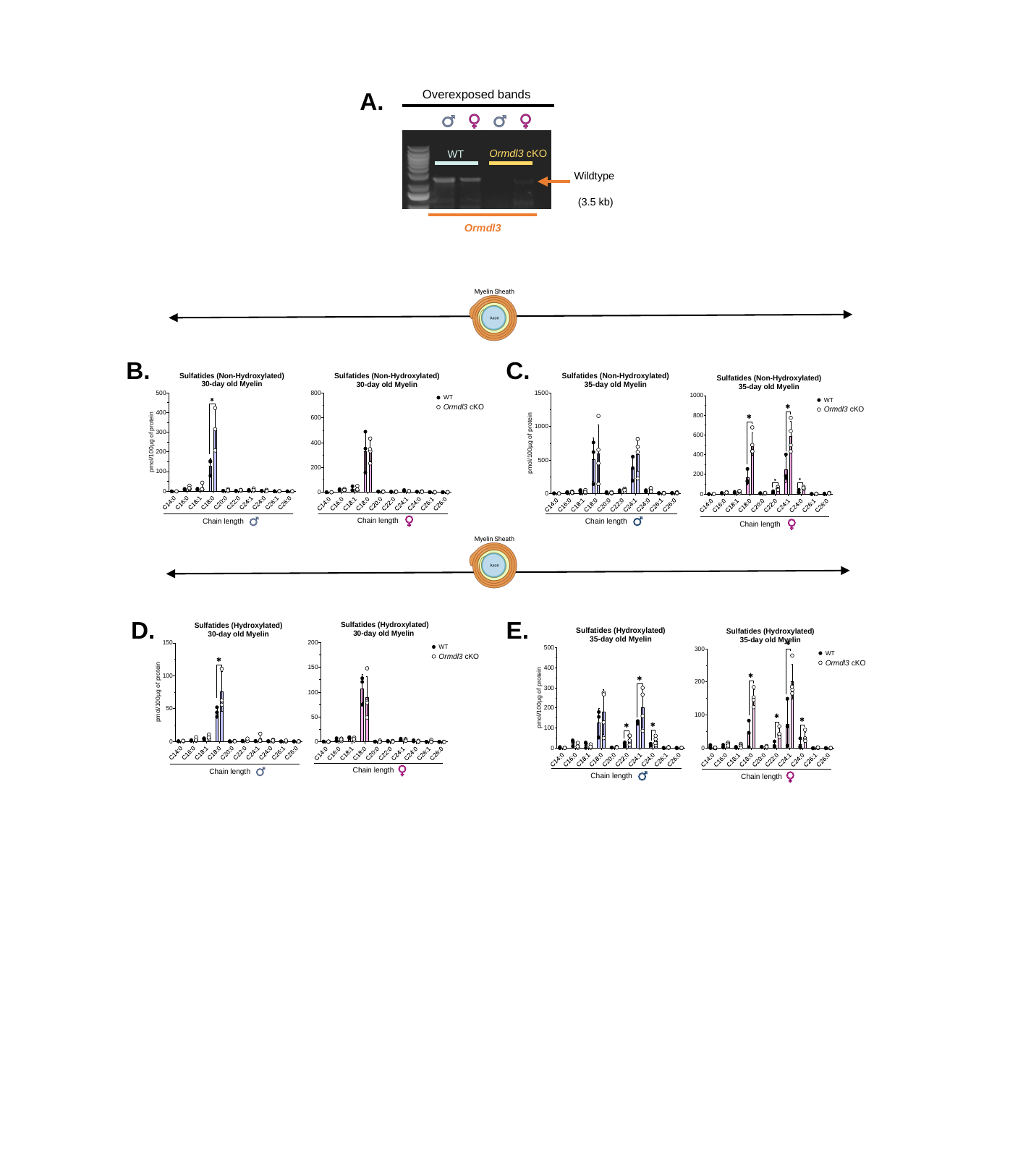

A.
Overexposed bands
Ormdl3 cKO
WT
Wildtype
(3.5 kb)
Ormdl3
B.
C.
D.
E.

### Slide 2
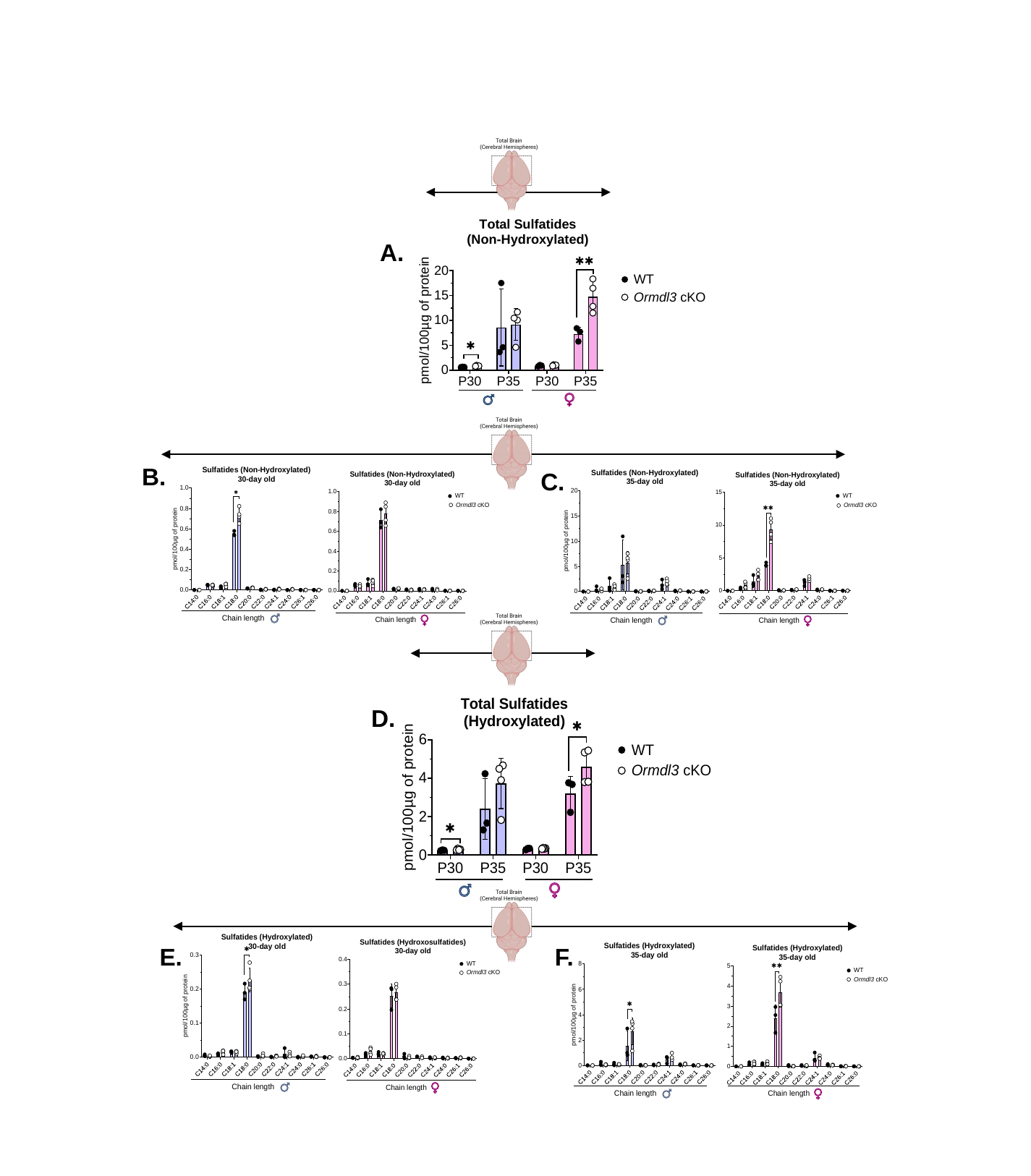

A.
B.
C.
D.
E.
F.

### Slide 3
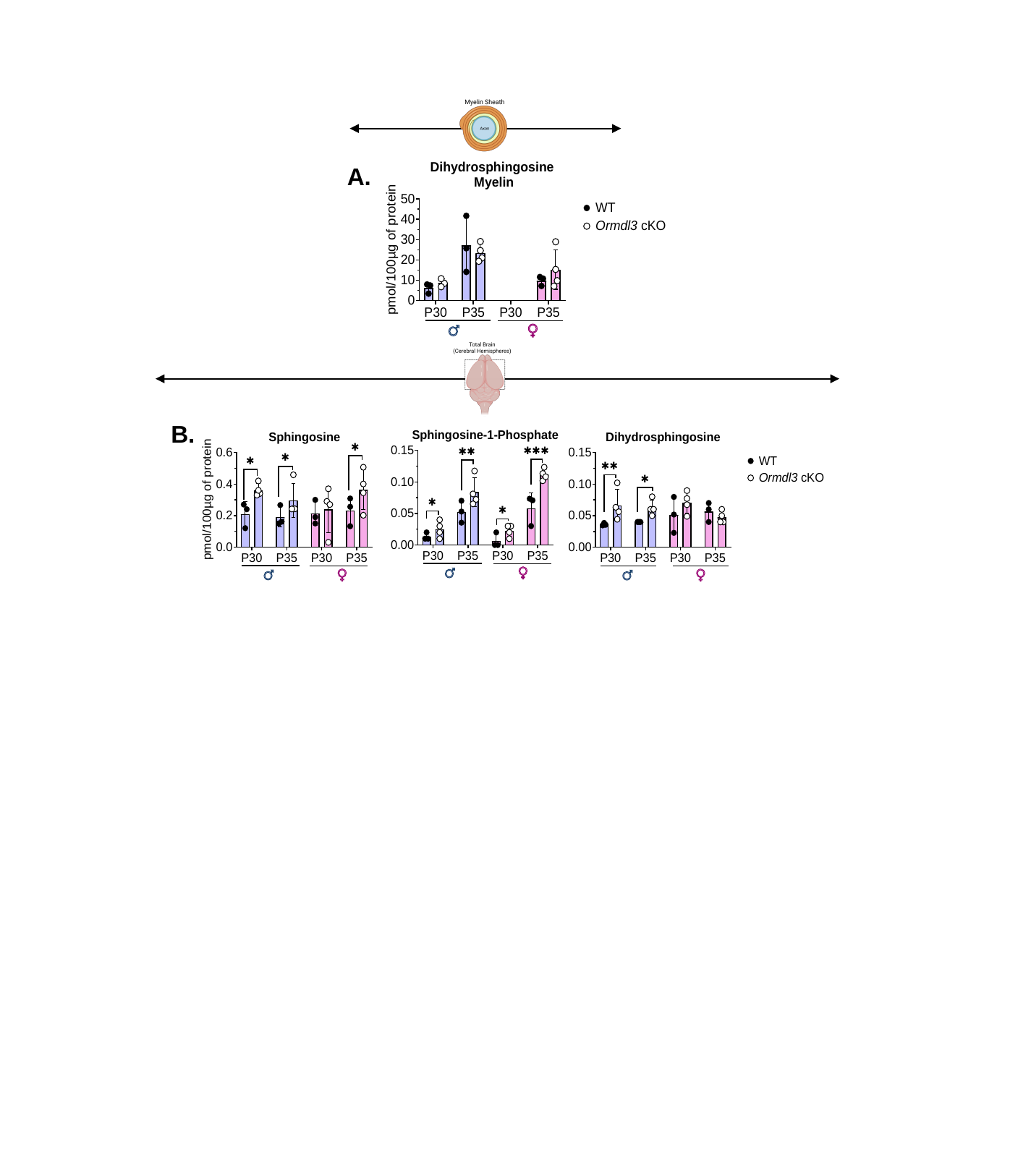

A.
B.

### Slide 4
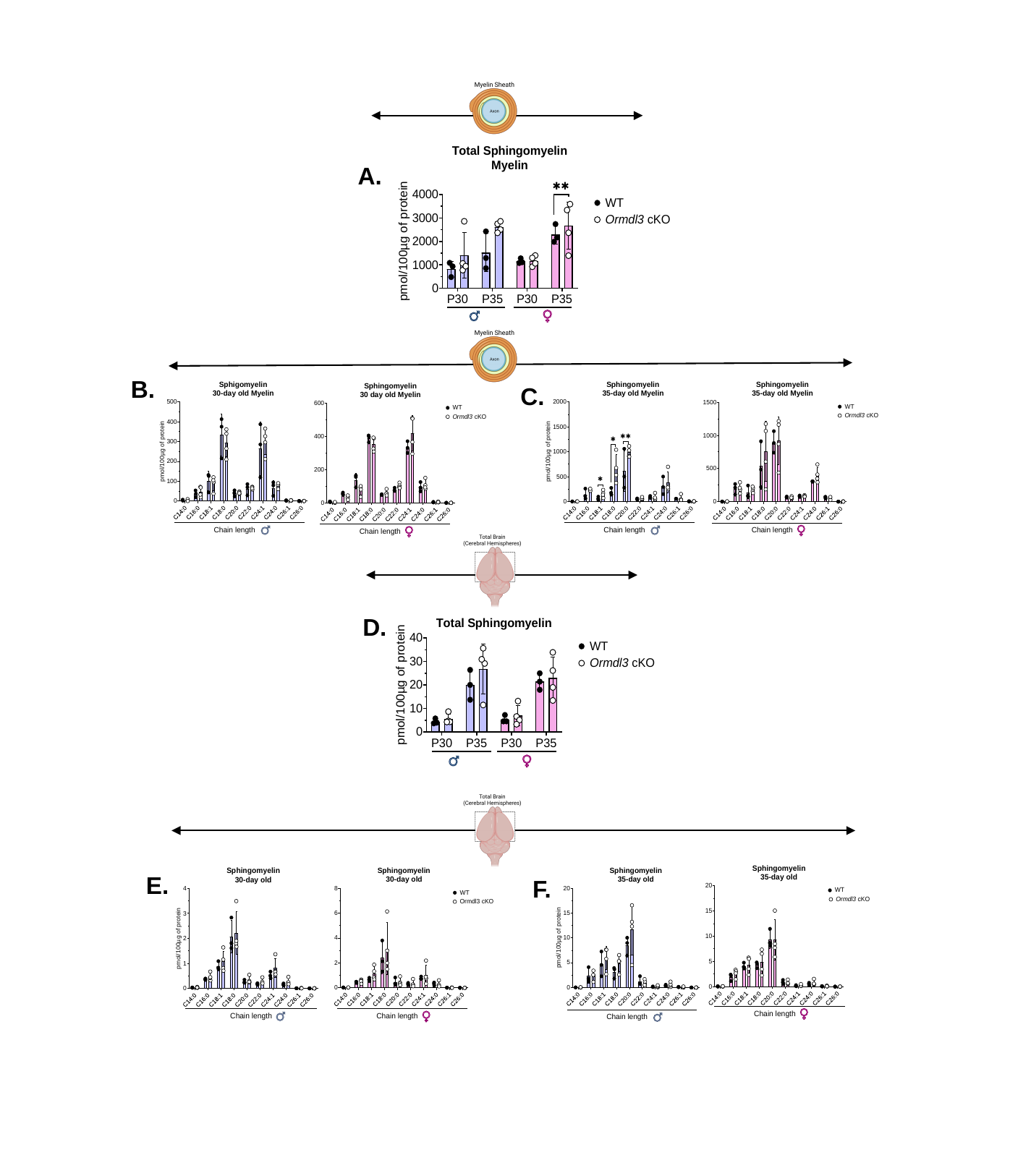

A.
B.
C.
D.
E.
F.

### Slide 5
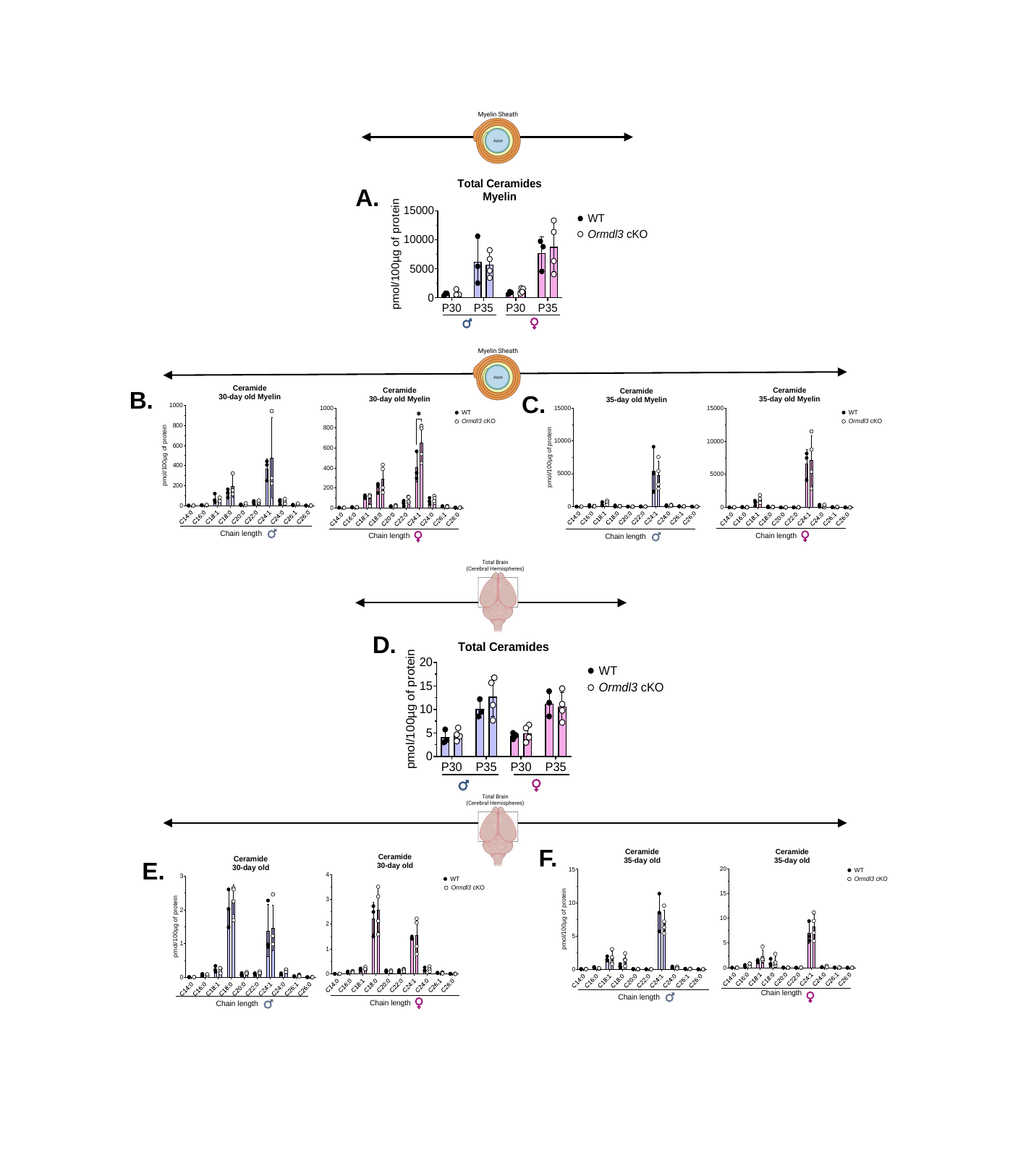

A.
B.
C.
D.
F.
E.

### Slide 6
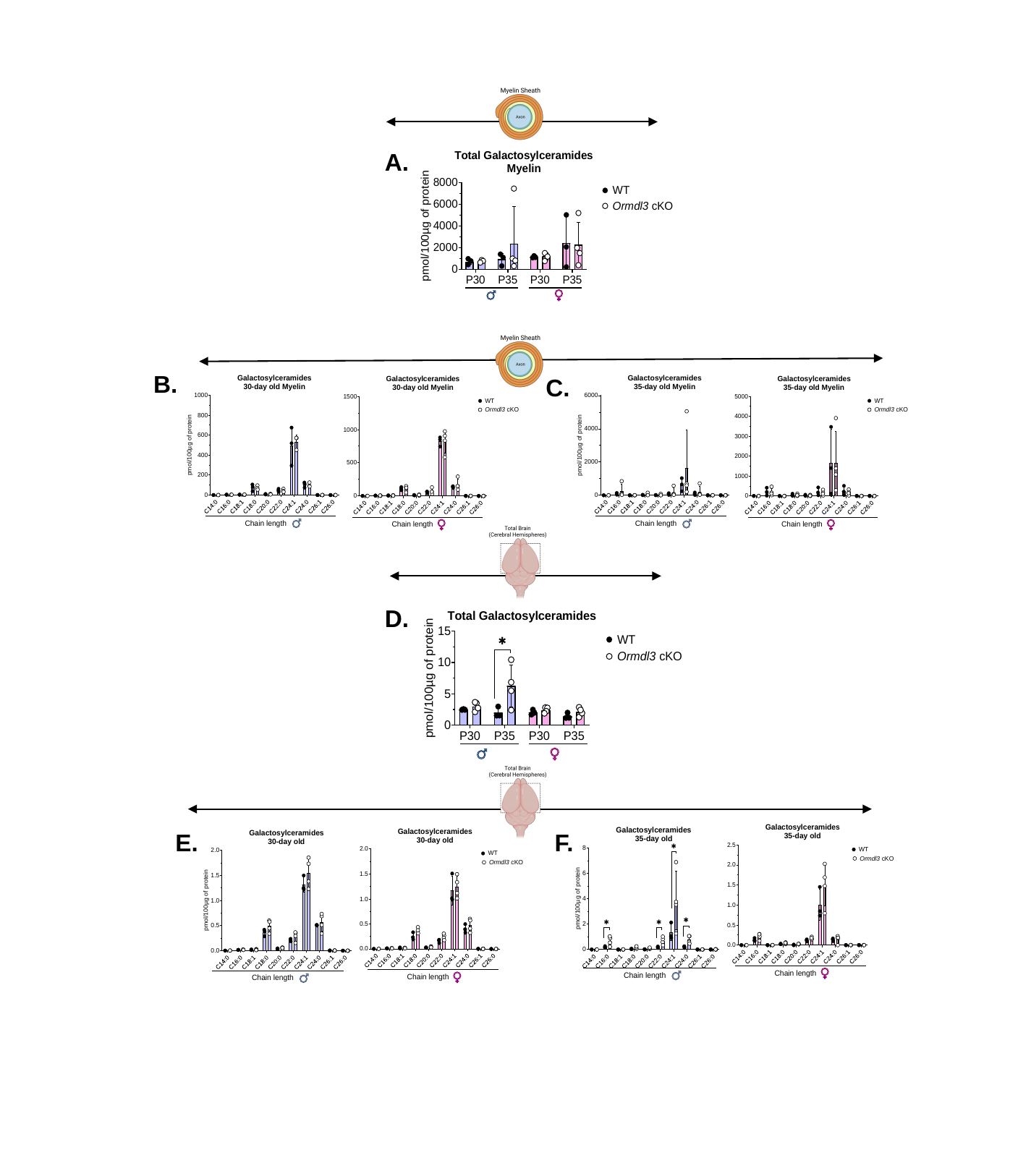

A.
B.
C.
D.
E.
F.

### Slide 7
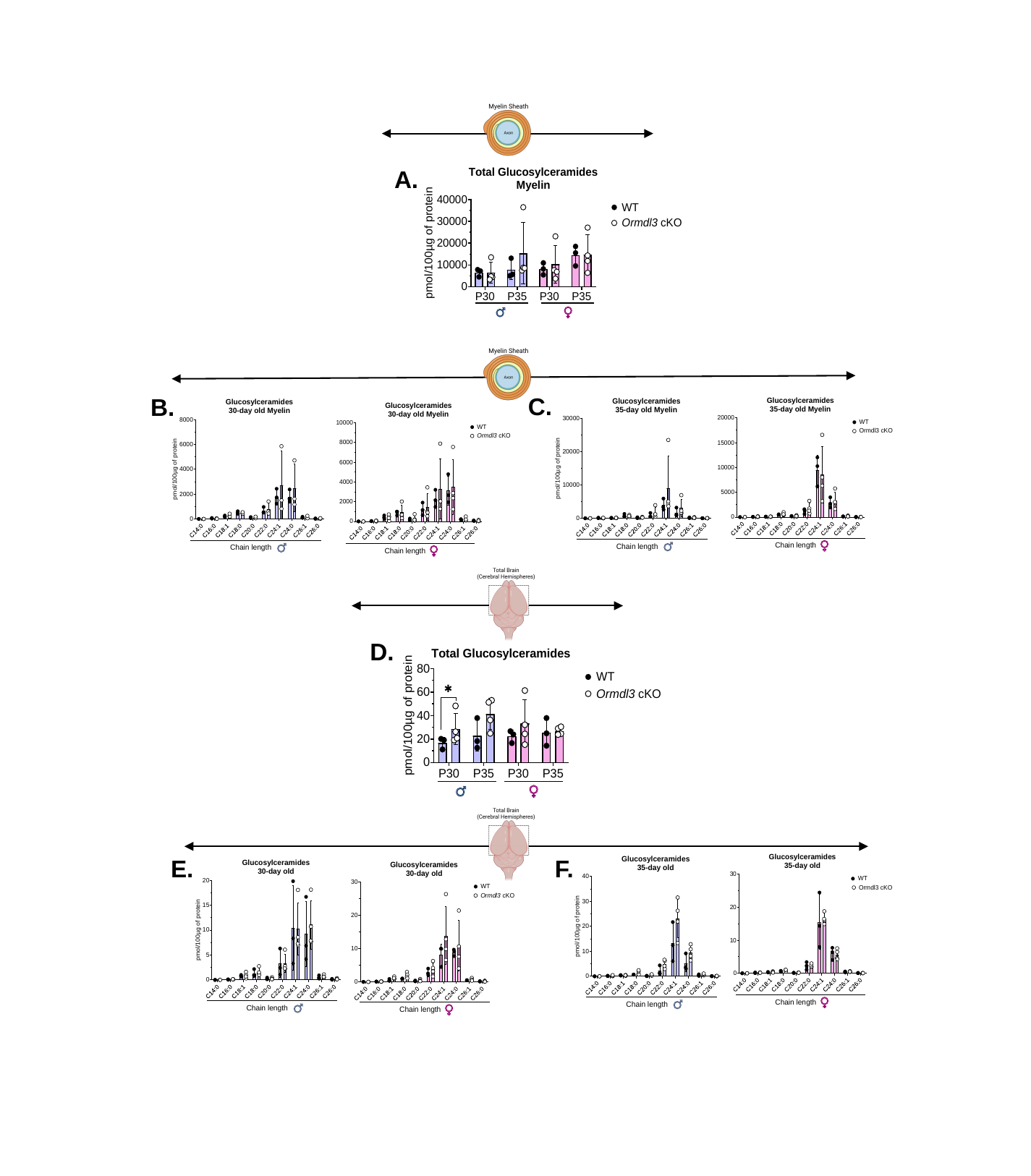

A.
C.
B.
D.
E.
F.

### Slide 8
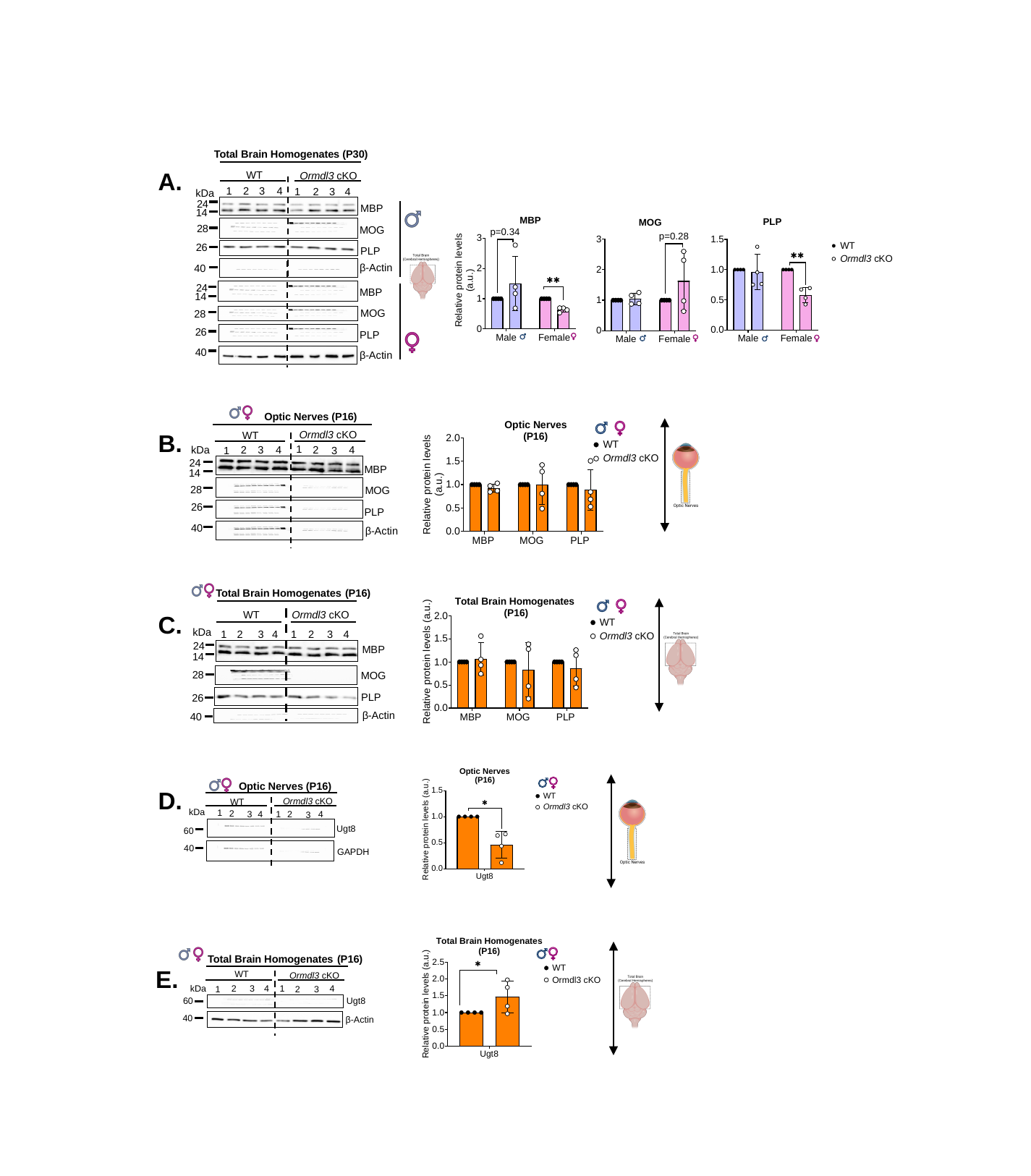

WT
Ormdl3 cKO
1
2
3
4
2
1
3
4
kDa
24
MBP
14
28
MOG
26
PLP
β-Actin
40
24
MBP
14
MOG
28
PLP
β-Actin
Total Brain Homogenates (P30)
A.
26
40
Ormdl3 cKO
WT
1
2
4
kDa
2
3
4
1
3
24
MBP
14
28
MOG
26
PLP
40
Optic Nerves (P16)
β-Actin
B.
Total Brain Homogenates (P16)
WT
kDa
2
1
2
3
4
1
3
4
24
MBP
14
28
MOG
PLP
26
Ormdl3 cKO
β-Actin
40
C.
Optic Nerves (P16)
Ormdl3 cKO
WT
kDa
1
2
3
4
1
4
2
3
Ugt8
60
40
GAPDH
D.
Total Brain Homogenates (P16)
WT
Ormdl3 cKO
3
kDa
1
4
2
4
2
3
1
Ugt8
60
40
β-Actin
E.
